# Developmental gene expression patterns driving species-specific cortical features

**DOI:** 10.1101/2025.02.18.638637

**Authors:** Awais Javed, Lucia Gomez, Veronica Pravata, Quentin Lo Giudice, Moein Sarhadi, Silvia Cappello, Esther Klingler, Denis Jabaudon

## Abstract

The cerebral cortex shows species-specific variations in size and organization, likely accounting for distinct behavioral abilities. These structural differences may reflect evolutionary changes in the developmental expression of shared genes. To investigate this possibility, we compared cell-type-specific gene expression across species in the developing mouse and human neocortex, and human cortical organoids, by generating a shared transcriptional reference framework. This identified genes with conserved/divergent expression patterns, providing a molecular foundation to interrogate species-specific cellular properties. Using this resource, we discovered that the transcription factor JUNB is expressed in human but not mouse progenitors. Through cell-type-specific gain- and loss-of-function experiments in mice and human organoids, we demonstrate that JUNB bidirectionally controls human cortical features, including progenitor proliferation rates, neuronal production timing, and total neuronal output. This reveals how cell-type-specific regulation of shared genes during development can drive species-specific cortical features, providing a framework for understanding the molecular basis of cortical evolution.

## Introduction

The neocortex, a hallmark of mammalian brains, is central for higher cognitive functions. It is composed of numerous populations of neurons that can be distinguished by several features, including their molecular identity, morphology, and connectivity, which ultimately determine their function (1–3). During neocortical development, glutamatergic neurons are born from progenitors called apical radial glia (RG), either directly, or through transit-amplifying cells called intermediate progenitors (IPs); deep-layer neurons are born first and preferentially project subcortically, while superficial-layer neurons are born later and mainly project intracortically (4–8). This process is largely conserved across mammals, but notable species-specific differences exist. For example, humans have a more diverse array of progenitors than mice, including basal and truncated RG, which are thought to contribute to cortical expansion and neuronal diversity (9–12). In addition, human neurons mature particularly slowly (13,14), arguably accounting for extended periods of environment-dependent plasticity. Several primate-specific genes influence cortical development: NOTCH2NL and ARHGAP11B enhance progenitor expansion, while SRGAP2 and LRRC37B regulate synaptic maturation timing (15–18). However, such species-specific genes are relatively rare, suggesting that temporal variations in the expression of shared genes, rather than species-specific genes, are a major source of interspecies diversity. Supporting this view, the timing of SATB2 expression controls whether interhemispheric axons traverse through the corpus callosum (in eutherians) or the anterior commissure (in marsupials) (19,20). Similarly, differences in the timing of expression of similar gene sets drive intracortical projections towards the motor or secondary somatosensory cortex (21,22). Hence, differences in the developmental expression timing of corresponding genes can lead to meaningful differences in brain features (23).

To systematically identify and compare cell-type-specific developmental gene expression dynamics during corticogenesis, we analyzed publicly available single-cell RNA sequencing datasets from mice (Mo), humans (Hu) and human organoids (Org_H_). We generated a common cell-type specific and temporal (*i.e.* "cyto-temporal") transcriptional framework allowing unbiased and systematic identification of conserved or divergent gene expression patterns. Our analysis reveals that while RG and immature neurons have highly conserved transcriptional programs, IPs display marked species-specific differences, particularly in genes controlling cell cycle properties and neurogenic output. This comparative framework further enables systematic identification of genes with divergent expression patterns between species and contexts, providing a valuable resource for understanding human-specific aspects of cortical development and for optimizing *in vitro* models of human brain development. Leveraging this dataset, we identify JUNB, a transcription factor with mutually exclusive expression in Mo neurons and Hu RG. Using gain- and loss-of-function experiments in Mo and Org_H_, respectively, we find that JUNB bidirectionally controls the developmental emergence of human features, including progenitor proliferation rates and neuronal production. Together, our results reveal how cyto-temporal gene expression regulation shapes cortical development across species and environments and shed light on the mechanisms underlying species-specific cortical features.

## Results

To normalize and compare developmental ortholog gene expression dynamics between species (*Mus musculus vs. Homo sapiens*) and contexts (*in vivo vs. in vitro*), we created a two-dimensional cellular manifold. Cortical neurons were positioned based on their gene expression along two orthogonal axes: normalized age (X-axis, from early to late corticogenesis), and differentiation state (Y axis, from RG to IP and glutamatergic neurons (N)) (**Fig. 1A**) (24,25). For each condition, we integrated several scRNAseq datasets from developmental stages where RG, IP and N were present and randomly selected balanced number of cells per cell type and age: embryonic day (E)12-E17 in Mo (4 datasets, 30’000 cells) (25–28), post-conception week (Pcw) 12-24 in Hu (3 datasets, 28’867 cells) (29–31), and 1-7 months *in vitro* for Org_H_ (3 datasets, 28’045 cells) (32–34), (**Fig. 1B**; **Fig. S1**; **Table 1**; **methods**). Original cell type annotations were curated into RG, IP and N after integration of the datasets based on unbiased clustering and expression of *SOX2*, *EOMES,* and *NEUROD6* (**Fig. S1**). Hu bRG clustered with aRG and were considered as RG for the analyses. Cell position within the manifold was determined using ordinal regression models predicting embryonic age (X-axis) and differentiation states (Y-axis) (**Fig. 1C**; **Figs. S2-4**; **methods**). This layout enables visualization of cyto-temporal gene expression as distinct genetic "landscapes" by color-coding cells based on gene expression levels (**Fig. 1D**). All data are available in an interactive format at www.humous.org (**Fig. 1E**; **Fig. S5**).

**Figure 1.**
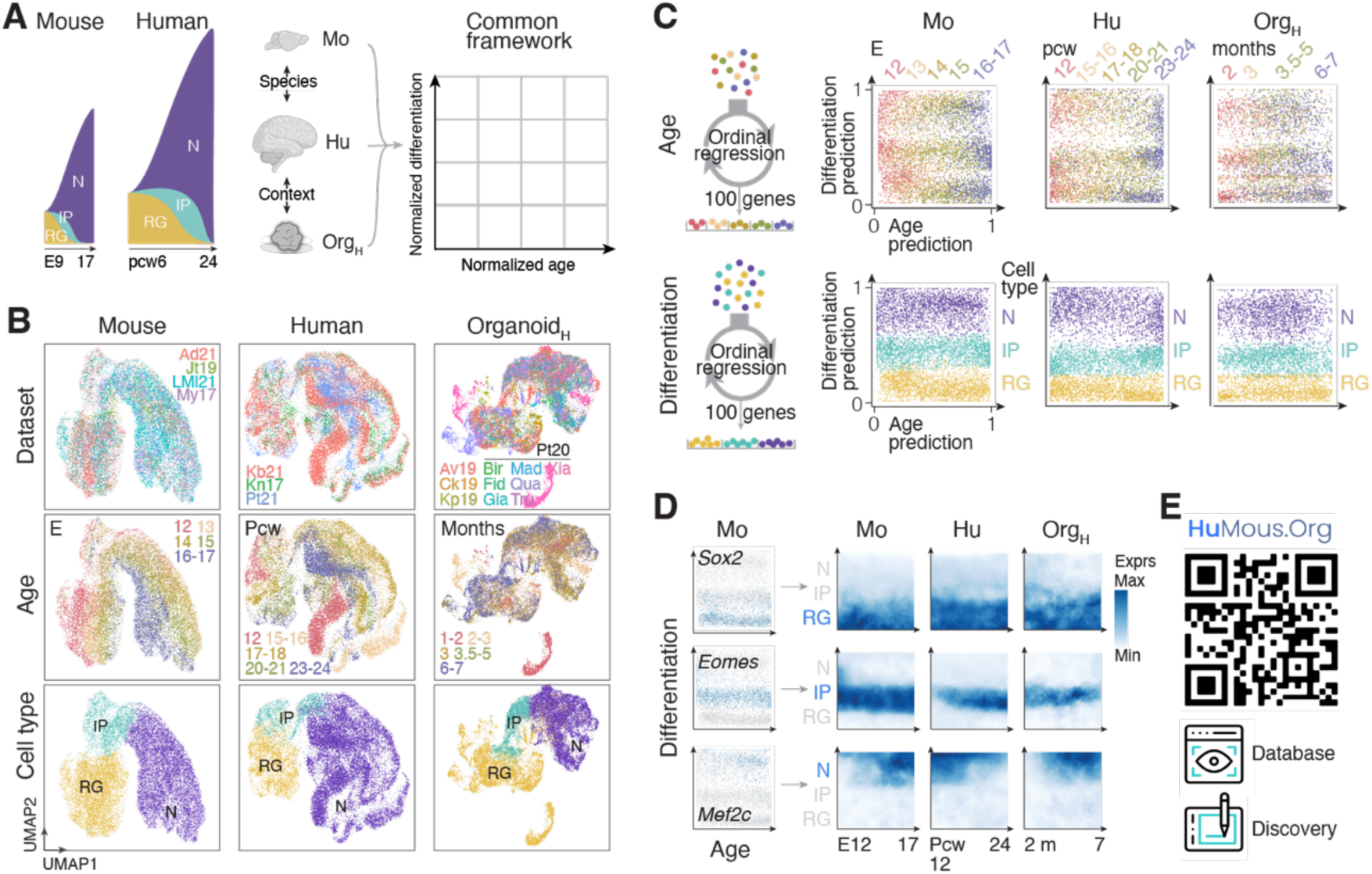
A common framework for comparing cell-type specific developmental gene expression across species and contexts. **(A)** Left: schematic representation of developmental cortical growth in mice and humans, with radial glia (RG), intermediate progenitors (IP) and neurons (N). Right: schematic representation of the two-dimensional cellular manifold used to analyze cyto-temporal gene expression patterns across species (Mo *vs.* Hu) and contexts (Hu *vs.* Org_H_). **(B)** UMAP representation of the datasets used in this study (top), age distribution of cells (center) and cell types (bottom). **(C)** Left: schematic representation of the ordinal regression model used to identify the top 100 genes driving embryonic age (top) and differentiation (bottom) identity. Right: Distribution of all cells from (B) using the framework presented in (A). **(D)** Representative gene expression landscapes showing how the manifold enables visualization of cyto-temporal patterns through color-coding of expression levels. **(E)** Link to the interactive web platform (www.humous.org) in which all data are interrogatable for database constitution and gene discovery. Abbreviations: UMAP, Uniform Manifold Approximation and Projection; Mo, mouse; Hu, human; Org_H_, human-derived cortical organoids; Pcw, postconception week; E, embryonic day.

As a first step in comparing landscapes, we determined their cyto-temporal organization using the standard deviation (SD) and entropy of expression as parameters. This distinguished three profiles of landscapes: non-patterned, ubiquitous, and patterned, (**Figs. S6A-C**; **S7A**; **methods**). Most landscapes (about 67.4%) did not display cyto-temporal patterning, *i.e.* gene expression was diffuse and patchy (low SD - low entropy). About 3.1% of landscapes had a ubiquitous profile, *i.e.* showed diffuse, homogenous expression (high SD - high entropy). Finally, patterned landscapes (29.5% of all landscapes) showed expression with the most obvious biological relevance, as gene expression was contrasted across cells and concentrated at specific times and/or differentiation states (high SD - low entropy) (**Fig. 2A**; **Figs. S6A,B**; **S7A**). Genes with ubiquitous landscapes were involved in generic cellular processes such as RNA processing and translation. In contrast, genes with patterned landscapes had ontologies related to specialized cellular functions, such as neuron migration and synapse organization (**Fig. S7B,C**). The fraction of patterned landscapes was significantly lower in Hu than Mo (*P* = 1.29 x 10^-126^; **Fig. 2A**; **Fig. S7**). This may reflect either increased pleiotropy in humans, where genes serve multiple functions across cell types and timepoints, and/or the presence of more human-specific isoforms that cannot be distinguished by the sequencing approaches used in these datasets. This differential patterning showed functional relevance, as assessed using gene ontology analysis: Hu-specific patterned genes were enriched for DNA repair and protein stability, supporting the extended maintenance of genome integrity during slower differentiation (35,36), while Mo-specific patterned genes related to ribosome biogenesis and synapse organization, potentially enabling rapid protein synthesis and neuronal maturation during a comparatively compressed development (**Fig. S7D,E**). Genes patterned in Hu but not Org_H_ were enriched for chromosome segregation, suggesting *in vitro* environmental influences on cell division (**Fig. S7F**) (37).

**Figure 2.**
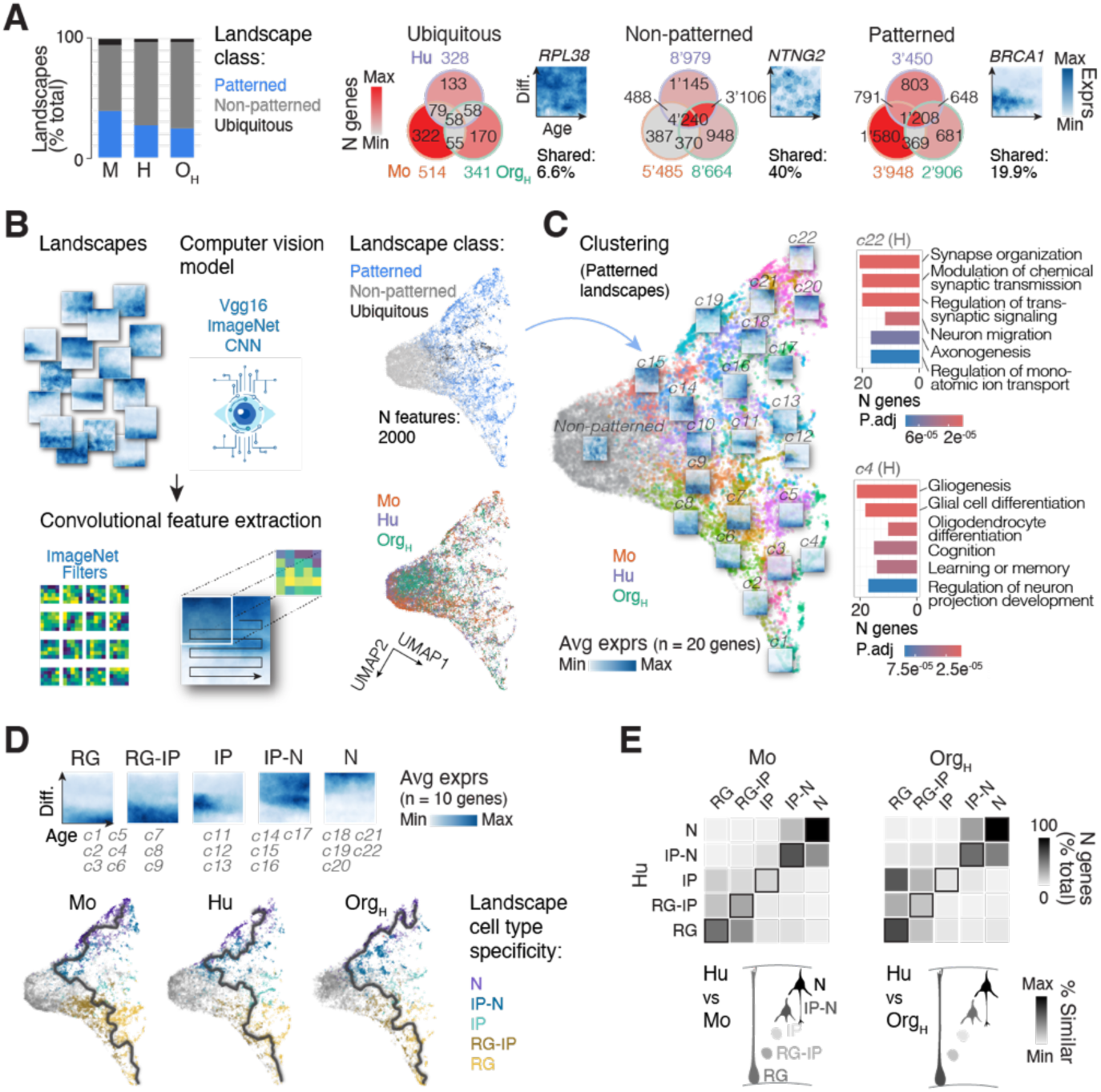
Systematic analysis of developmental gene expression patterns reveals cell-type specific conservation. **(A)** Classification of gene expression landscapes based on standard deviation and entropy metrics, revealing three main profiles: non-patterned (67.4%), ubiquitous (3.1%), and patterned (29.5%). **(B)** Left: schematic representation of the landscape analytic pipeline. Landscapes were clustered based on 2,000 computer vision-extracted features. Right: UMAP visualization of landscape distributions. Each dot represents a single landscape. **(C)** Left: Organization of the 21 identified landscape clusters. Right: Example landscapes from key clusters showing associated biological processes (Human data). **(D)** Top: Landscape clusters categorized based on cell-type specific expression patterns. Bottom: Pseudotime alignment using Monocle showing a bottom-to-top organization of differentiation in the landscape UMAP. **(E)** Top: Cross correlation analysis demonstrating conservation of gene expression patterns in RG and N populations but divergence in IP across species (Mo *vs.* Hu, left) and contexts (Hu *vs.* Org_H_, right). Bottom: schematic representation of the data shown in the cross correlations. Abbreviations: RG, radial glia; IP, intermediate progenitor; N, neurons; Avg exprs, average expression.

We focused our attention on genes with patterned landscapes given their straightforward biological relevance. As a first approach to analyze their diversity, we applied a computer vision approach (38) to extract 2’000 variable features of pixel distribution per landscape. These features were used as parameters to display landscapes in a Uniform Manifold Approximation and Projection (UMAP) space (**Fig. 2B**; **Figs. S6C**; **S8**; **methods**). The resulting visualization revealed 21 clusters, each representing distinct cyto-temporal patterns of gene expression (**Fig. 2C**). These clusters were organized along a differentiation axis, starting from landscapes showing specific expression in RG (clusters 1 to 7), and ending with N-specific landscapes (clusters 18 to 22) in all three conditions (**Fig. 2C**, left; **Table 2**). Genes involved in sequential cellular processes during corticogenesis, such as genes with ontologies related to cell cycle, neuron migration, projection development and synapse development and function, were sequentially organized along this axis (**Fig. S9A**; **methods**). For examples, in Hu, cluster 4 was enriched in genes involved in gliogenesis (*e.g. BCAN*, *GFAP*), cluster 18 contained genes involved in dendrite morphogenesis and axonogenesis (*e.g. CDK5*, *SEMA3A*), and cluster 22 genes involved in synapse organization (*e.g. CAMK2B*, *FLRT2*) (**Fig. 2C**, right; **Table 2**), consistent with differences in progenitor fate progression (from neuron- to glia-producing), neuron development, and functional maturation, respectively.

As a systematic approach to understanding transcriptional variation across differentiation, we first analyzed gene distribution patterns across key cell types (RG, IP, and N), and then examined how these patterns evolved over developmental time. Starting with cell types – most clusters showed cell type specificity rather than temporal specificity (**Fig. S8C,D**) – we defined five categories of landscape based on cellular expression: RG-only, RG and IP, IP-only, IP and N, and N-only (**Fig. 2D**, top). IP-only landscapes were more frequent in Hu than Mo and Org_H_, suggesting specialized molecular controls over this population compared to Mo, where IPs were less prominent, and Org_H_, where IPs are scarce and poorly characterized (39) (**Fig. 2D**, bottom; **Fig. S8B**). RG-only and N-only expression patterns showed high conservation across species and contexts, while IP patterns diverged markedly (**Fig. 2E**; **Fig. S8**), suggesting that developmental transcriptional start and end points within this time window are evolutionarily stable, while intermediate progenitor identities are more variable.

Expression comparison of genes defined by their developmental ontologies revealed distinct organizational principles across species. In Mo, developmental genes follow a strict sequential expression pattern: cell-cycle genes are expressed in RG/IP, followed by migration genes in IP/N, and finally projection/synapse genes in N. In contrast, Hu shows more overlapping patterns, with IPs maintaining expression of cell-cycle genes while RGs simultaneously express migration and projection-related genes (**Fig. S9A,B**). Additionally, correlation analysis revealed that the degree of conservation between species varies by cell type: ontologies of genes expressed in progenitors (RG and early IPs) showed low correlation, while ontologies of genes expressed in neurons showed high correlation (**Fig. S9C**). Together, these findings reveal two distinct aspects of species-specific cortical development: different temporal organization of developmental programs and divergent evolution of progenitor-specific gene expression.

We next compared temporal gene expression dynamics between species by classifying landscapes based on relative cyto-temporal expression. This revealed sets of genes with either delayed (n = 288) or precocious (n = 176) expression in Hu *vs.* Mo (**Fig. S10A**). In RGs, delayed Hu genes were enriched for cell division ontologies (*e.g. NUF2*, *PLK1*), while precocious ones related to protein folding (*e.g. CRYAB*, *PPIC*). In IP-Ns, genes delayed in Hu were linked to synapse function (*e.g. NRP1*), while precocious ones related to axon development and cytoskeleton organization (*e.g. SEMA6D, STMN2*) (**Fig. S10B,C**; **Table 3**). This cyto-temporal uncoupling - earlier axon development programs but delayed synaptic maturation genes - may reflect a key feature of human cortical development in which basic wiring is poised early on to detect environmental signals that then sculpt synaptic connections during prolonged periods of development (18,40–43).

To identify genes whose expression diverged most across species and context, we next measured pairwise correlation values between landscapes patterned in all conditions (*i.e.* Mo, Hu, and Org_H_; **Fig. 3A**; **methods**). This analysis identified four classes of landscapes: (i) conserved, with high correlation across all conditions; (ii) species-specific, *i.e.* differing between Mo and Hu but conserved between Hu and Org_H_; (iii) context-specific, *i.e.* differing between Hu and Org_H_ but conserved across species, potentially highlighting environment-sensitive genes; and (iv) divergent, *i.e.* showing low correlation for all comparisons. Genes with conserved expression patterns across all conditions formed the largest category (26%), while only 8% were condition-specific (**Fig. 3A**), indicating broad conservation of gene regulatory programs. The divergent expression patterns we observed were not explained by basic genomic or protein features – neither by chromosomal location, coding sequence conservation, peptide length, nor functional class (**Fig. 3B**; **Fig. S11**). Instead, these patterns reflected specific biological functions: conserved landscapes were enriched for genes involved in fundamental neuronal processes like synapse organization (*e.g. SYT4*, a key regulator of neurotransmitter release). Species-specific landscapes were enriched for genes controlling more specialized features such as axon fasciculation (*e.g. CNR1*, which guides axon organization), while context-specific landscapes showed enrichment for cell division genes (*e.g. CENPA*, essential for chromosome segregation) (**Fig. 3C-E**; **Table 4**). Together, these findings demonstrate that a majority of gene expression programs are preserved across species and contexts, with some specific to either species or environmental conditions.

**Figure 3.**
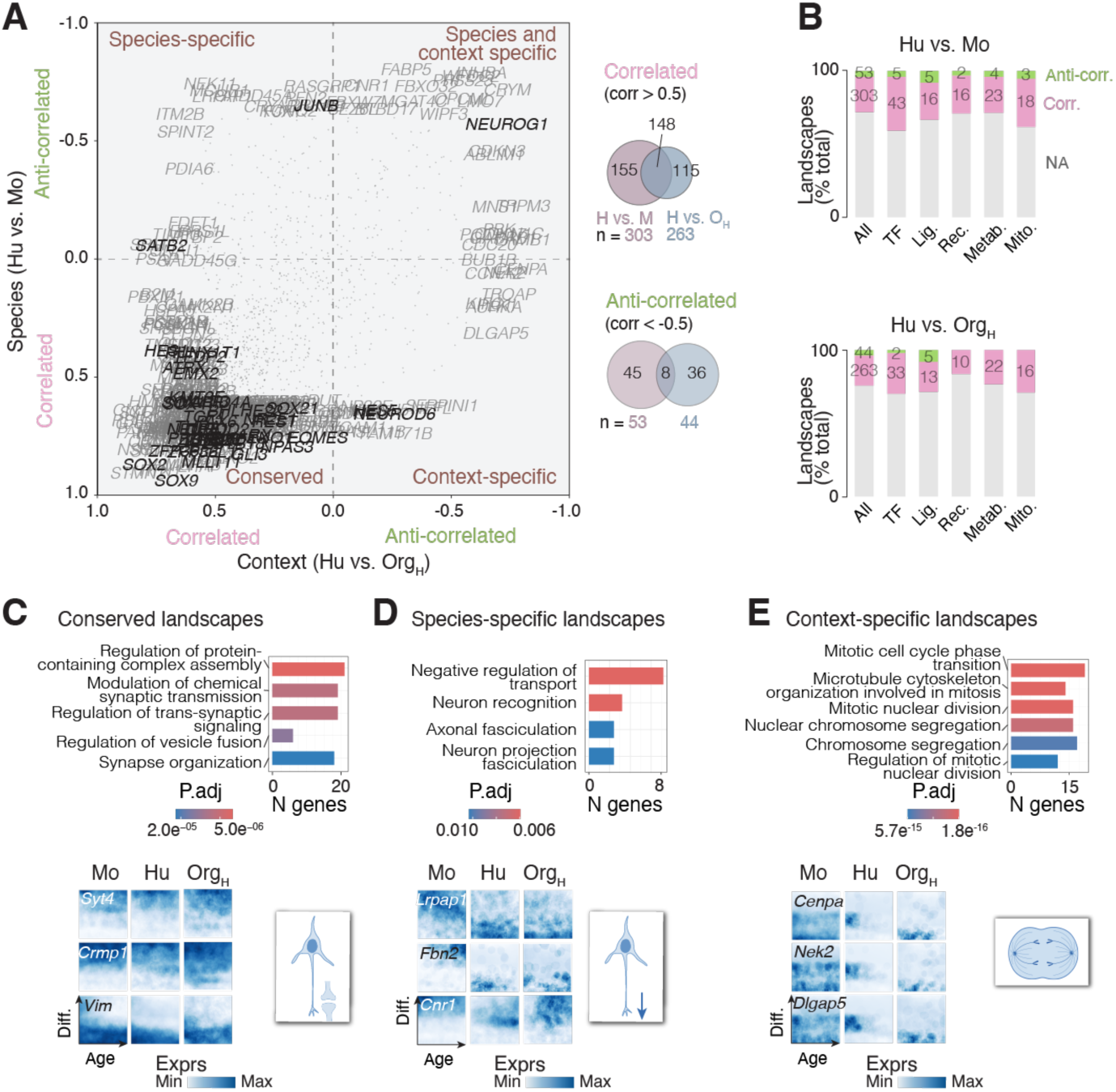
Identification of conserved and divergent gene expression programs across species and contexts. **(A)** Left: Pairwise correlation analysis of gene expression landscapes across species (Hu *vs.* Mo, Y axis) and context (Hu *vs.* Org_H_, X axis), revealing four classes of genes: Conserved, Species-specific, Context-specific, and Divergent landscape patterns. Transcription factors are in black. Right: overlap in correlated (top) and anti-correlated (bottom) landscapes in species and context comparisons. **(B)** Distribution of correlated and anti-correlated landscapes between species (top) and context (bottom) for select gene classes. TF: transcription factors; Lig.: ligands; Rec.: receptors; Metab: metabolic; Mito.: mitochondrial. **(C-E)** Representative landscapes and enriched biological processes for conserved (*e.g. SYT4*), species-specific (*e.g. CNR1*), and context-specific (*e.g. CENPA*) genes. Abbreviation: Mo, mouse; Hu, human; Org_H_, human-derived cortical organoids; NA, not assigned; Corr., correlated; Anti-corr., anti-correlated; P.adj, adjusted p-value; Diff, differentiation; Exprs, normalized expression.

To investigate the functional impact of divergent expression patterns, we searched for genes that showed strong species-specific differences. Among highly anti-correlated genes *in vivo*, *JUNB*, a component of the AP-1 transcription factor complex that regulates cellular responses to growth and stress signals (44–47) was the transcription factor exhibiting the most anti-correlated expression pattern (corr. = −0.64; **Fig. 3A**). *JUNB* is composed of one exon showing 86.5% similarity between Hu and Mo (96.7% between Hu and macaque; **Fig. S12A**). In Mo, *Junb* is expressed in N, while in Hu and Org_H_, it is restricted to RG (**Fig. 4A**; **Fig. S12B**). In macaque-derived cortical organoids, *JUNB* is also enriched in RG and absent in N (**Fig. S12B)** (48). This is in contrast with other members of the AP-1 complex, whose cyto-temporal expression is mostly conserved (**Fig. S12C**). Immunostaining confirmed *JUNB*’s cell-type specificity, with protein expression restricted to progenitors lining the ventricle in Org_H_ and no expression in Mo VZ (**Fig. 4A**).

**Figure 4.**
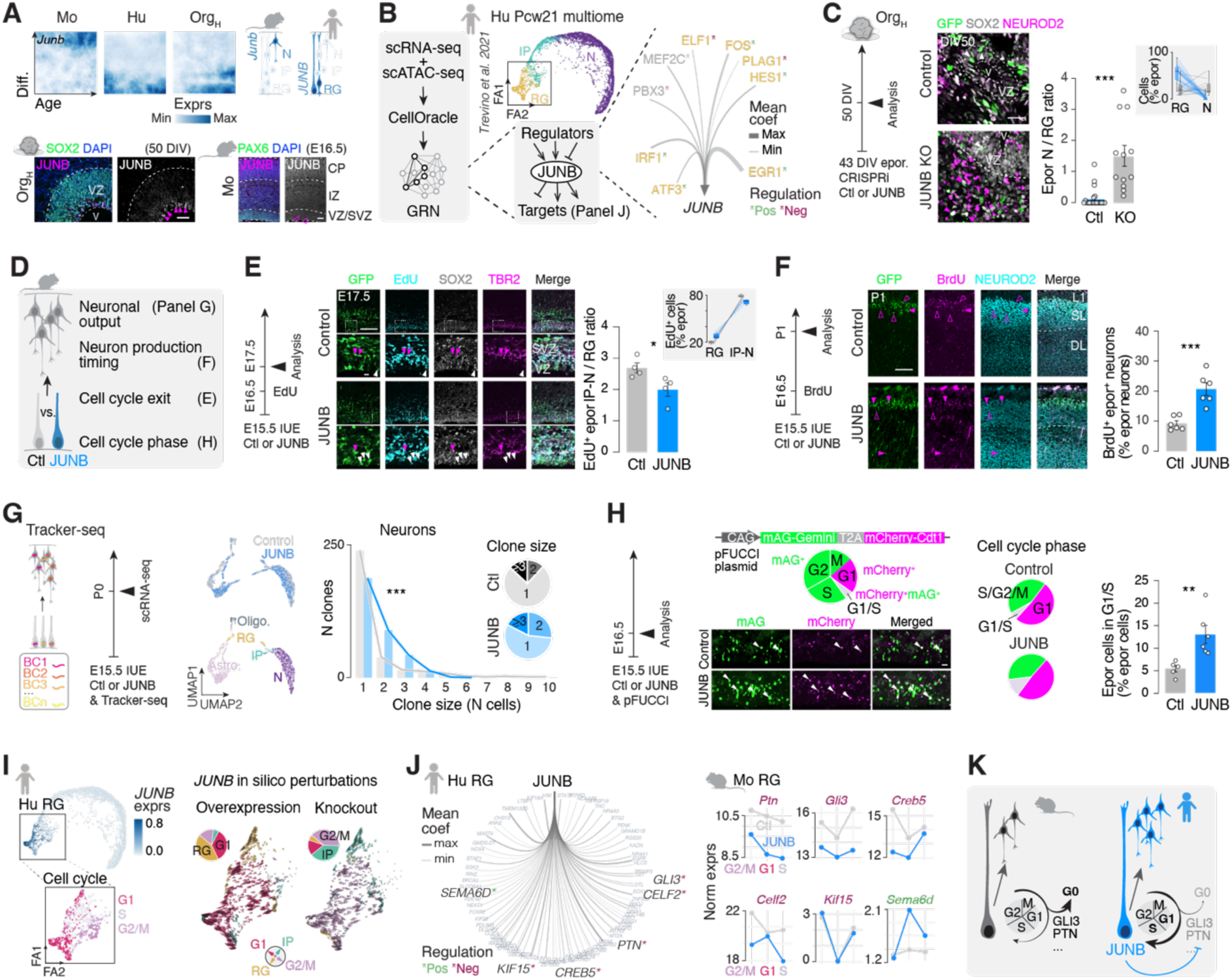
JUNB bidirectionally controls species-specific cortical progenitor properties. **(A)** Top left: Landscapes for *JUNB* showing specific expression in Hu RG. Top right: schematic representation of cell-type specific expression. Bottom: Immunostaining validation of JUNB in the ventricular zone of Org_H_ and Mo. Magenta arrowheads show JUNB^+^ progenitors in Org_H_, empty arrowheads show JUNB^-^ progenitors in Mo. **(B)** Left: Schematic of the approach used to identify JUNB gene regulatory network (GRN) in Hu RG. Center: Human multiome data (31) cell types. Right: Chord diagram representing the predicted upstream regulators of *JUNB*. The width of the lines connecting to *JUNB* shows the scaled value of mean coefficient score calculated in CellOracle GRN analysis. Factors expressed in Hu RG are shown in yellow. Positive (green) and negative (bordeaux) regulations are indicated. **(C)** CRISPRi-mediated JUNB knockout in human-derived cerebral organoids and neuron-to-progenitor ratio. White arrowheads, electroporated SOX2^+^ progenitors; empty magenta arrowheads, non-electroporated NEUROD^+^ neurons, filled magenta arrowheads, electroporated NEUROD2^+^ neurons; each dot representing a ventricle (n = 4-5 organoids / condition). Inset showing raw values for RG and N proportions of electroporated cells per ventricle. **(D)** Summary of JUNB functional characterization in Mo. **(E)** Left: summary of the Mo JUNB acute overexpression experimental design. Center: representative photomicrographs following immunocytochemistry. Right: Quantification of E16-born (EdU-positive) electroporated IP-N to RG ratio at E17.5. White arrowheads, EdU^+^/SOX2^+^/ TBR2^-^ RG; magenta arrowheads, EdU^+^/TBR2^+^ IP. Inset shows raw values for RG and IP-N proportions of electroporated cells per embryo. **(F)** Left: summary of the Mo JUNB long-term overexpression experimental design. Center: representative photomicrographs following immunocytochemistry. Right: Quantification of E16-born (BrdU-positive) electroporated N at P1. **(G)** Left: Tracker-seq principle and summary of the clonal lineage tracing experimental design. Center: UMAP showing distribution of JUNB-overexpressing cells (top) and cell types (bottom). Right: Clonal analysis revealing the number of neurons per clone born from control or JUNB-expressing RG. Pie charts summarizing clone size proportions in the two conditions. **(H)** Left: Summary of the pFUCCI reporter experimental design. Center top: schematic representation of the plasmid construct and correlation of cell-cycle phase with color. Cells express both colors (*i.e.* white cells) only during the G1-S transition. Center bottom: Representative photomicrographs following immunocytochemistry. Arrowheads, mCherry^+^/mAG^+^ RG in the VZ. Right: Pie charts showing the proportions of electroporated RG in the different cell cycle phases and quantification of electroporated RG in G1/S phase in both conditions. **(I)** Left: Expression of *JUNB* in Hu single-cell multiome data (top) and RG cell cycle phase annotation (bottom). Right: Prediction of Hu single cell trajectories upon *JUNB* overexpression or knockout using CellOracle *in silico* analysis. Pie charts representing the proportions of trajectories towards each cardinal direction (*i.e.* G1, G2M, RG, IP). **(J)** Left: Chord diagram representing predicted downstream targets of Hu JUNB dysregulated upon JUNB expression in Mo RG. The width of the lines shows the scaled value of mean coefficient score calculated in GRN CellOracle analysis. Positive (green) and negative (bordeaux) regulations are indicated. Right: Expression of differentially expressed Hu JUNB targets in control and JUNB-overexpressing Mo RG. Targets predicted to be negatively regulated by JUNB in Hu are in bordeaux, and targets predicted to be positively regulated are in green. **(K)** Summary of the findings on how JUNB controls species-specific cortical progenitor properties leading to increased neuronal output. Abbreviations: VZ, ventricular zone; v, ventricle; DIV, days *in vitro*; E, embryonic day; P, postnatal day; Pcw, postconception week; Norm exprs, normalized expression; Diff., differentiation; Mean coef, mean coefficient; Avg exprs, average expression; IUE, *in utero* electroporation; Epor, electroporated; Ctl, control; RG, radial glia; IP, intermediate progenitor; N, neuron; SL, superficial layers; DL, deep layers; BC, barcode; FA, factor analysis; Oligo, oligodendrocyte, Astro, astrocyte. Scale bars: (A Org_H_, C) 25 μm; (A, Mo) 50 μm; (E,F) 100 μm; (H) 20 μm.

To identify transcriptional regulators that could account for RG-specific JUNB expression in humans, we applied CellOracle analysis (49) to a human multiome dataset (31) (Pcw21). This approach identified several RG-specific transcription factors as potential upstream regulators of *JUNB* - including FOS, EGR1, ATF3 and IRF1 - which showed lower / absent expression in mouse RG (**Fig. 4B**; **Fig. S12D**). Remarkably, despite their absence in mice, the binding motifs for these regulators were present and accessible in both Mo (E14-E16) and Hu (PCW21) *JUNB* promoters, as shown by motif enrichment analysis (50) (**Fig. S12E**, **Table 5**). This suggests that *JUNB* promoter is poised for activation in mouse RG but remains inactive without these human-specific regulators.

To examine if this pattern of conserved binding sites but species-specific regulation extends beyond *JUNB*, we analyzed the relationship between gene expression patterns and motif conservation across species for all expressed genes. However, we found no correlation between how similarly genes are expressed in Mo versus Hu and how conserved their binding motifs are at the promoter (**Fig. S12F**). This suggests that differences in the combination of regulators present and potential enhancers, rather than binding site evolution at the promoter, may underlie species-specific gene expression. Finally, to determine whether the *JUNB* upstream regulators identified above coordinate a broader species-specific program in RG, we analyzed their predicted targets using CellOracle. We found that these transcription factors also regulate other human RG-specific genes like *GLIS3* and *CEBPD* (**Fig. S12D**), suggesting that JUNB is part of a broader human gene regulatory network in RG.

To examine the functional effects of JUNB’s RG-specific expression pattern in Hu, we performed gain- and loss-of-function experiments. We started with *JUNB* knockdown using CRISPRi in Org_H_, which led to an increase in neuron-to-progenitor ratio, consistent with accelerated neurogenesis (**Fig. 4C**; **Fig. S13A**). This suggests that JUNB participates in maintaining the prolonged proliferative state of human RG. Supporting this interpretation, expressing JUNB in mouse RG using *in utero* electroporation at E15.5 caused an increase in RG numbers at the expense of IP, consistent with an increase in RG self-replication (**Fig. 4D,E**; **Fig. S13B**). Accordingly, neuronal generation was extended, as demonstrated by a higher fraction of neurons born after the first 24 hours post-electroporation (**Fig. 4F**). Finally, to examine whether this shift in RG self-replication ultimately led to an increase in neuronal production, we performed clonal analysis using Tracker-seq (51), which revealed larger clone size in neurons born from JUNB-expressing RG (**Fig. 4G**; **Fig. S13C,D**). Together, these bidirectional manipulations demonstrate that JUNB is both necessary and sufficient to promote the extended proliferative capacity characteristic of human RGs

To investigate how JUNB regulates RG self-replication, we co-electroporated JUNB with the cell-cycle reporter pFUCCI (52). This experiment revealed an increased proportion of JUNB-expressing RGs in G1/S transition (mCherry^+^/mAG^+^ cells in VZ/SVZ; **Fig. 4H**; **Fig. S13E**), suggesting that JUNB promotes G1-to-S (proliferative) over G1-to-G0 (differentiative) transitions. In support of this interpretation, single-cell RNA sequencing of mouse RGs performed one day after *in utero* electroporation showed an enrichment in G1/S phase cells (**Fig. S13F,G**). Accordingly, *in silico* manipulation of JUNB levels in human cortical multiome data using CellOracle revealed that JUNB knockdown shifted RGs away from a G1 phase signature towards an IP fate, while overexpression enhanced G1 phase RG features (**Fig. 4I**; **methods**). Together these data demonstrate that JUNB controls RG proliferation through regulation of cell cycle progression.

We next sought to identify the downstream molecular mechanisms involved. Analysis of the human JUNB regulatory network identified 81 putative target genes, of which six showed congruent and significant changes in our mouse scRNAseq data (5 downregulated and 1 upregulated; **Table 6**). These six genes showed the highest differential expression during G1/S phase, consistent with the results above (**Fig. 4J**; **Fig. S13H**). *PTN*, *GLI3*, and *CREB5* decreased during human but not mouse corticogenesis, while *KIF15* was expressed in mouse but absent in human RGs (**Fig. S13H**). Notably, GLI3 and PTN are known to promote neuronal differentiation over RG proliferation, including through lengthening of G1 phase (53,54), suggesting that JUNB acts as a molecular regulator between mouse-like and human-like RG properties through repression of these differentiation regulators (**Fig. 4K**).

## Discussion

The results presented here provide a comprehensive insight into cortical cell-type specific and temporal patterns of gene expression across species (mouse and human) and contexts (*in vivo* and *in vitro*). Through systematic analysis of developmental transcriptional landscapes, we reveal that evolutionary conservation varies markedly across cell types - while radial glia and neurons show relatively conserved expression patterns, intermediate progenitors exhibit striking divergence in gene expression. This finding supports a model of mosaic brain evolution (55,56), where distinct cell populations evolve at different rates. The high divergence in intermediate progenitors suggests they may be key drivers of interspecies differences in cortical architecture through species-specific controls of neuronal output.

Our functional analysis of JUNB illustrates how temporal regulation of shared genes can drive species-specific features. JUNB’s ability to induce human-like progenitor properties when expressed in mouse tissue suggests the presence of latent transcriptional programs that can be activated in the right cyto-temporal context. This represents a parsimonious evolutionary mechanism where subtle changes in cyto-temporal expression of conserved genes can produce dramatic phenotypic differences - similar to how minor timing changes in circadian regulators can broadly impact metabolism and physiology (57). The shift in neuronal production timing in JUNB-expressing RGs could account for a key human feature: disproportionate expansion of superficial cortical layers. Since these layers primarily contain neurons that connect different cortical areas, this seemingly subtle timing change can profoundly impact cognitive capabilities by altering the balance between intracortical – *i.e.* associative – and subcortical communication.

While our study focused on excitatory neurons, similar principles likely apply to inhibitory interneurons. Given their extended developmental trajectories and heightened plasticity, even subtle interspecies differences in gene expression timing could significantly impact their circuit integration and function. This may be particularly relevant for understanding human-specific aspects of cortical inhibition and its disorders.

The comparison between human tissue and organoids revealed both strengths and limitations of *in vitro* models. While many developmental programs were well preserved, we observed differences in cell cycle-related genes, suggesting that aspects of progenitor division may require signals present in vivo, such as factors in cerebrospinal fluid (58). Looking ahead, it will be important to understand the upstream mechanisms controlling these temporal expression differences, including epigenetic regulation, metabolic state, and post-transcriptional control (59–61). Building on the framework established here, such investigations will further illuminate how genetic programs, temporal regulation, and environmental factors interact to generate species-specific brain features during development and evolution.

## Methods

### Data source and selection

All single-cell transcriptomics data used in ^2,4–8^ are open public datasets from mouse (25–28) and human (29–31) embryonic cortices, and human derived cortical organoids (32–34) (**Table 1**). Gene expression datasets were filtered for cortical excitatory lineage cells across the corticogenesis period (*i.e.* post-conception weeks (Pcw)12 to 24 for human, embryonic days (E)12 to 17 for mouse, and 4 to 37 weeks for organoid_H_). We used authors’ original cell annotations for filtering cell types corresponding to RG, IP, as well as immature and mature neurons and only selected ages at which both RG, IP and neurons were detected.

### Data integration and cell type annotation

We integrated datasets belonging to the same condition to alleviate batch effects. For mouse datasets, we randomly selected 6’000 cells per age group, defined as E12, E13, E14, E15 and E16-17. For human datasets, we randomly selected 6’000 cells per age group, defined as Pcw12, Pcw14-16, Pcw17-18, Pcw20-21, and Pcw23-24, except for Pcw12, where only 4’867 cells were available. We then used the Seurat V3 integration pipeline (62) for pairwise dataset integration over the 2’000 most variable features across datasets (*VST* method). We specified 15 *CCA* dimensions and 20 neighbors for anchoring and regressing out the effect of the total number of genes detected per cell (*nFeature_RNA*). For organoid datasets, due to partially overlapping group definitions in original data annotations and imbalanced data availability, we aimed at obtaining the same number of cells as for other species (∼30’000 cells) and defined flexible cell selection windows per stage *in vitro*, as follow: 1-2 months: 5’706 cells, 2-3 months: 6’294 cells, 3 months: 8’644 cells, 3.5-5 months: 599 cells, and 6-7 months: 8761 cells. Pairwise integration was referenced to the previously integrated human gene expression matrix (see **Fig. S1**) and organoid integration units were defined as the combination of dataset and protocol to avoid capturing experimental biases in organoid generation.

Each integration object was used for defining differentiation groups consistently across datasets. To do so, we calculated 2D UMAP reductions from integrated gene expression matrices and used clustering for delimiting differentiation groups into RG, IP and neurons, based on *SOX2*, *EOMES* and *NEUROD2* marker expression respectively, resulting in 2’425 cell per differentiation group in mouse, 2’450 in human and 1’800 in organoid_H_ (**Fig. S1**).

### Ordinal modelling for age and differentiation

To convert qualitative age and differentiation groups into gene expression aware quantitative continua, we used a custom adaptation of SVM-based ordinal regression (bmrm R package, https://github.com/pradosj/bmrm) on integrated data for each condition. For ordinal model training we selected a balanced number of cells, n = 50 per age and differentiation groups (*e.g.* 250 cells total in mouse) - the biases due to datasets or protocols were balanced by the integration, see above. Two sequential ordinal trainings were performed: the first (full), on all scaled genes and the second (reduced) on the model’s top 50 scaled genes as weighed ranking of the full model (25 for each extreme of the ordinal output). Full and reduced ordinal model training followed a cost matrix defined on the levels of the factor to predict (age or differentiation groups). Twenty cross validations were implemented for reduced model training.

To homogenize age and differentiation axes reconstruction across conditions, we rescaled the resulting ordinal score from 0 to 1. Additionally, cell distribution across ordinal scores were normalized towards unimodality and uniformity, using a custom function for distribution detection, tahn transformation, and outlier correction. First, the unimodality and uniformity of ordinal score vectors’ distribution was assessed using dip (dip.test R function, diptest package) and Kolmogorov Smirnoff testing (ks.test R function, stat R package), respectively. Unimodal and uniform scores were left unchanged. Non-uniform scores followed a smoothing transformation applied to each ordinal target level (age or differentiation groups) and consisting of applying the hyperbolic tangent function (tanh, base R package) over the median of the sub-score with a strength adapted to its standard deviation. Ordinal normalization output was winsorized for outlier correction (substitution of outlier values by closest within the quantile window of retention: 0.01 and 0.99 probabilities, Winsorize R function, DescTools package) and rescaled on a 0 to 1 range for homogenization.

### Gene expression landscapes

#### Data filtering

Before landscape generation, we excluded genes expressed in less than 1% of the cells across datasets within a condition. As a result, we selected for calculating gene landscapes *n* = 12’757, *n* = 12’714, and *n* = 13’564 genes for human, mouse and organoid_H_ datasets, respectively. Expression for all these genes was normalized and scaled regressing out the total number of genes detected per cell (*nFeature_RNA*).

#### Landscape generation

For each condition independently, we used scaled RNA-seq data and applied the following procedure: we first constructed a KNN array (k nearest neighbors, k=100) using the cells’ age and differentiation ordinal scores as neighborhood coordinates. Next, for each gene, we averaged its expression across blocks of 100 neighbor cells to construct a smoothed expression array, so-called gene landscape, of dimension 250 x 250 (averaged cells, *i.e.* pixels), as we have done before (24,25). Each gene landscape was stored for further analysis as an array and as a png image (using a custom R function that combines the raster and ggplot2 R libraries, png dimensions were set at 244 x 244 pixels for optimal functioning of computer vision modelling, see below).

### Definition of gene expression landscape profiles based on entropy and standard deviation

The entropy and standard deviation were calculated for each gene landscape using the entropy R function from the entropy library and the sd R function, respectively. The standard deviation (SD) was calculated on the gene landscape vectorized array (as it is a measure of distribution), while the entropy (E) was calculated on the image matrix (to measure the level of randomness in pixel patterning). Landscape expression classes were defined as follow: E < 20 and SD < 0.1 = “low”; 20 < E < 25: 0.1 < SD < 0.15 = “non-patterned”, SD > 0.15 = “patterned”; E > 25 and 0.15 < SD < 0.35 = “ubiquitous”. Genes with low expression profiles were expressed by very few cells sparsely located on the age-differentiation axes, genes with non-patterned profiles by many cells distributed in a random manner on age-differentiation axes, genes with ubiquitous profiles by many cells homogeneously located on age-differentiation axes, while genes with patterned profiles were expressed by few to numerous cells but with a clear pattern on age-differentiation axes (see **Fig. S6**).

### Landscape computer vision feature extraction

For extracting spatially relevant features of gene landscapes, we encoded them using a computer vision model, VGG16, pre-trained on the ImageNet dataset, accessed using the keras API through an R miniconda with python version 3.7. Specifically, we cut the sequential model layers to retrieve the output from the 5^th^ max pooling layer, providing 25’088 features per landscape. Gene landscapes’ 244 x 244 pngs were preprocessed for VGG16 prediction to be converted to arrays of shape [1,7,7,512]. Next, we launched the model prediction on each image and flattened the output to obtain a feature vector (1 x 25’088), all finally bind in a data matrix ready for clustering.

### Gene landscape clustering based on computer vision features

Gene landscapes of the three conditions were compiled into a Seurat object using Seurat R package (v5.1.0), with the assay containing their 25’088 VGG16-extracted features, and the metadata all information about each landscape, including their entropy and SD. Iterative clustering was next used to cluster the landscapes. For this, we normalized the features and defined the 2’000 most variable using the FindVariableFeatures function. Data were scaled and the principal components (PC) calculated using these variable features. Clustering iterations were performed using 1:15 PC. The first clustering iteration (resolution = 0.2) allowed to identify landscapes with low, non-patterned, expression profile. After excluding these landscapes, the variable features were redefined, as well as the principal components, neighbors and clusters (resolution = 0.5). This second clustering iteration allowed to label two more clusters with non-patterned expression profiles. Following the same strategy, we excluded these landscapes and made a third clustering iteration (resolution = 1.3) using only the patterned landscapes (defined based on entropy and standard deviation, see above), which resulted in 22 clusters of gene expression landscapes.

#### Annotation of expression landscapes

Clusters of gene expression landscapes were manually annotated, based on their age (early, late, non-specific) and cell type (RG, RG-IP, RG-N, IP, IP-N, N, non-specific) specificities, by averaging 20 random landscapes per cluster using a standard base R mean of the array of landscapes. For age and type annotation illustration in **Fig. 2**, 50 landscapes per annotation were averaged using the same procedure.

### Pseudotime trajectories of gene expression landscapes

To build each trajectory, the numerical representation of patterned landscapes in the UMAP space were considered and used as input for R package Monocle3 workflow (V1.4.20). The previously described Seurat object was converted into a cell_data_set objet with the help of SeuratWrappers (V0.3.5) package and processed with the classical Monocle3 Trajectories workflow (https://cole-trapnell-lab.github.io/monocle3/docs/trajectories/). The cluster_cells function was used with the k=10 and reduction_method=”UMAP” parameters, the learn_graph function with use_partition=F and minimal_branch_len=3 parameters, and the order_cells function with default parameters. The Choose_graph_segment function was used to connect starting and ending nodes, respectively corresponding to the less and most advanced branching point of the pseudo-differentiation tree. Only the main trajectory per condition is shown.

### Landscape correlations

Aiming at obtaining a comparable numerical metric of landscape similarity, we calculated correlations between gene landscape and all others across intra- and inter-species. We defined a custom R function taking the 0-1 normalized vectorial representation of two landscapes and thresholding it at 0.3-pixel intensity value for diminishing the impact of noise on correlation estimates. As correlation is impacted by sample size, a robust implementation was set where for each pair of landscapes to compare, we bootstrapped the calculation 10 times for a fixed set of 2000 pixels (sampling with replacement) equally placed on the two landscapes. Finally, we calculated the median across the 10 bootstrapped correlations. Gene landscapes with a bootstrapped correlation coefficient higher than 0.5 across conditions were tagged as highly correlated, while landscapes with correlation smaller than −0.5 were tagged as anti-correlated.

### Gene lists

To look at genes with specific functions, we used published databases of mammalian metabolic enzymes (63) and mitochondrial proteome (64) (MitoCarta3.0), SingleCellSignalR for ligands-receptors (65), and Transcription factors (66).

Genes with given ontologies were analyzed using ontologyIndex R package from the ontologyX suite (67).

### Humous.org website

An interactive web platform (**Fig. 1E**) was made available to serve both as a user-friendly landscape database (*browser* and *gene sets*), and to incorporate additional functionality to guide discovery for further research (*correlations* and *draw your landscape*). A summarized version of computational methods and data sources is as well included (*methods*). www.humous.org was created combining a JavaScript front-end and an R back-end, containerized in a docker network and hosted at University of Geneva servers. The *browser* page allows to search a gene (or series of genes) of interest and retrieve the respective landscapes across species. A tabular display enables a friendly layout for landscape comparison. The *gene sets* page behaves similarly but encodes a-priori defined groups of genes based on function or specificity. The *correlations* page enables users to recursively find genes that are similar or dissimilar within and across species, therefore guiding the discovery of candidate genes for further research (see landscape correlations methods section). The *draw your landscape* page encodes a landscape correlation-based engine for retrieving the top best landscape matches based on a user input drawing.

### Gene ontology analyses

Gene ontology over-representation analysis was performed for biological processes using the enrichGO function from the clusterProfiler package, with the following parameters: pAdjustMethod = “BH”, pvalueCutoff = 0.05, qvalueCutoff = 0.05, minGSSize = 10, maxGSSize = 500) (68).While top ontologies are shown in figures for readability, significant ontologies are in **Tables 3-4**.

### Plasmids

pCAG-GFP-ITR vector was a generous gift from Christian Mayer, which was modified to either include H2B sequence upstream to GFP (pCAG-H2B-GFP-ITR) or TurboRFP-NLS instead of the GFP (pCAG-TurboRFP-NLS-ITR). Addgene #29687 pCS2-Flag-JUNB was used for JUNB overexpression experiments. Addgene #175573 pX458-Ef1a-dCas9-KRAB-MECP2-H2B-GFP (CRISPRi) was modified by replacing the Ef1a promoter with CAG using Neb HiFi DNA assembly reaction (Neb: #E2621S). pCAG-CRISPRi plasmid generated in this study was then digested with BbsI-HF (Neb: #R3539S) and using Neb HiFi DNA assembly, gRNA1: GAGGCCAGCCTCGGAGCCAGC and gRNA2: GCCAGCTCCCTGCTGGCTCCG was cloned into the backbone. pCAG-CRISPRi without a gRNA was used as a control. Addgene #175266 pCAG-FUCCI was used for cell cycle phase analysis experiments.

### iPSCs lines and maintenance

iPSCs (HPS00076:409B2 and GM05399) were cultured on Matrigel (Corning) coated plates (Thermo Fisher, Waltham, MA, USA) in mTesR1 basic medium supplemented with 1x mTesR1 supplement (STEMCELL Technologies, Vancouver, Canada) at 37 °C, 5% CO_2_, and ambient oxygen level. Passaging was done by Gentle Cell Dissociation Reagent (STEMCELL Technologies) treatment.

### Generation, culture and electroporation of cerebral organoids

Cerebral organoids were generated from induced pluripotent stem cells (iPSCs) at ∼80% confluency according to previously published protocols (69,70). Cells were dissociated with Accutase and resuspended in low bFGF medium with 50 μM ROCK inhibitor. Live cells (2,000 per well) were seeded into low-attachment 96-well plates, forming embryoid bodies (EBs) after 24 hours. EBs were cultured for 5–6 days and subsequently transferred to Neural Induction Medium and cultured for 4–5 days. Neuroepithelial tissues formed, showing radially organized pseudo-stratified epithelium. On day 11, aggregates were embedded in Matrigel and cultured further. By day 18, organoids were placed on an orbital shaker at 57 rpm, with medium changes every 3-4 days, supporting cerebral tissue development. For electroporation of cerebral organoid ventricles, organoids with well-formed ventricles were selected at day 43 as previously published (71,72). Prior to electroporation, plasmid DNA was prepared at a concentration of 1-2 μg/μL in sterile nuclease-free water, with 0.05% Fast Green added to visualize injection. Microcapillaries were pulled from borosilicate glass using a Narishige PC-10 micropipette puller with one-step pulling at a heater level of 62°C and 4 weight blocks. The capillaries were loaded with the plasmid solution via capillary action. Each organoid was transferred individually into an electroporation dish containing pre-warmed (23°C) Opti-MEM. Using a stereomicroscope within a horizontal flow cabinet, microcapillaries were used to inject 5 μL of plasmid solution into the ventricles of each organoid, targeting specific lobes of interest. After injection, electroporation was performed using a BTX Harvard apparatus at 80 V with five pulses of 1 ms each and 1-second intervals. Following electroporation, the organoids were transferred to fresh medium and allowed to recover before further processing 7 days post electroporation.

### Mouse strains

CD1 pregnant female mice from Charles River Laboratory were used. The experimental procedures described here were conducted in accordance with the Swiss laws and previously approved by the Geneva Cantonal Veterinary Authority. All mice were housed in the institutional animal facility under standard 12 h:12 h light:dark cycles with food and water ad libitum.

### *In utero* electroporation of mouse embryos

*In utero* electroporations were performed as described in (73). Timed pregnant CD1 mice were anaesthetized with isoflurane (5% induction, 2.5% during the surgery) and treated with the analgesic Temgesic (Reckitt Benckiser). Embryos were injected in the lateral ventricle with ∼1 μL of DNA plasmid solution (diluted in endotoxin-free TE buffer and 0.002% Fast Green FCF (Sigma)). Embryos were then electroporated by holding each head between circular tweezer-electrodes (5 mm diameter, Sonidel) across the uterine wall, while 5 electric pulses (50 V, 50 ms at 1 Hz) were delivered with a square-wave electroporator (Nepa Gene, Sonidel).

### BrdU/EdU injection

On average, 350 μl of 10 mg/ml BrdU (ThermoFisher: #B23151) or 400 μl of 7.5 mg/ml EdU (ThermoFisher: #A10044) resuspended in 0.9% NaCl was intraperitoneally (IP) injected into pregnant females corresponding to 200 mg/kg (10 μl per g bodyweight).

### Tissue processing and immunohistochemistry

#### Organoid

For immunohistochemistry (IHC), organoid 20 µm thick sections were prepared on slides, which were thawed and rehydrated by rinsing in phosphate-buffered saline (PBS) for 5 minutes. The sections were blocked and permeabilized using a solution of 0.25% Triton-X, 150 mM glycine, and 4% normal goat serum (NGS) in PBS for 1 hour at room temperature in a humidified chamber. The following primary antibodies were diluted in blocking solution (4% NGS, 0.1% Triton-X in PBS) and applied to the slides overnight at 4°C: monoclonal mouse anti-SOX2 (1:500; AB_2665892; Protein Atlas, #AMAb91307), chicken anti-GFP (1:1000; Aves Labs, AB_10000240, #GFP-1020), monoclonal rabbit anti-NEUROD2 (1:300; Abcam, AB_10866309, #ab109406), rabbit anti-JUNB (1:100; Proteintech, AB_2129996, #10486-1-AP). After washing three times in PBS with 0.1% Tween for 5 minutes each, corresponding secondary antibodies from ChromoTek (alpaca anti-Mouse IgG1 nanobody: A488: sms1AF488-1, AB_2827578; A568: sms1AF568-1, AB_2827578; A647: smsG1CL647-1, AB_2941312) and ThermoFisher Scientific (goat anti-rabbit IgG (H+L): A546: A-11010, AB_2534077; A647: A-21244, AB_2535812; goat anti-chicken IgY (H+L) A488: A32931TR, AB_2866499) were diluted at 1:500 and applied for 1-2 hours at room temperature in the dark. A final wash in PBS without Tween was performed, and the slides were mounted for imaging with AquaPoly Mount.

#### Mice

Pregnant female mice were sacrificed by cervical dislocation and the embryos were removed from the abdominal cavity and placed in 1X HBSS for dissection. The embryos were first decapitated and transferred to a second petri dish in fresh 1X HBSS before the brain was dissected out. The electroporated cortices were screened for GFP and brains were placed in freshly prepared 4% PFA overnight at 4°C. P1 pups were injected with Esconarkon and perfused with 1X PBS followed by 4% PFA. Brains were then dissected out from the head and kept overnight in 4% PFA at 4°C.

The next day, the samples were washed with 1X PBS and 20% sucrose in PBS was added to cryoprotect the samples. The samples were then embedded in OCT and frozen on dry ice. Finally, the tissue was cut in 20 μm coronal sections using Leica Cryostat 3050S and mounted on Superfrost plus adhesion slides, which were kept at −20°C. Control and hJUNB electroporated samples were always mounted on the same slides and immunostained together. The slides were then washed with 1X PBS three times and placed in blocking buffer containing 3% BSA, 0.05% Triton in 1X PBS for one hour. The following primary antibodies were added to the slides in fresh blocking buffer and incubated overnight at RT: rabbit anti-SOX2 (1:200; ab97959; Abcam), rat anti-EOMES (1:200; Dan11mag; Invitrogen), rabbit anti-NEUROD2 (1:400; ab104430; Abcam), chicken anti-GFP (1:1000; ab13970, Abcam), rat anti-BrdU (1:500; ab6326, Abcam), rabbit anti-JUNB (1:50; 10486-AP1, Proteintech), monoclonal mouse anti-PAX6 (1:200, 13B10-1A10; ThermoFisher Scientific). After three washings with 1X PBS, the corresponding secondary antibodies were added at RT for 2 hours: goat anti-rabbit A555 (A-11008), goat anti-chicken A488 (A-11039), goat anti-rat A555 (A-21434). goat anti-rat 647 (A-21247). All secondary antibodies were from ThermoFisher and diluted at 1:500 in 1X PBS. DAPI was used as a counterstain at the end of the immunostaining protocol and slides were mounted with coverslips using Fluoromount-G mounting medium (00-4958-02, ThermoFisher).

EdU was revealed using a custom protocol as previously described (74) at the end of the immunostaining protocol. BrdU was revealed using a previously described protocol (75) in which slides were incubated with primary antibodies including anti-BrdU overnight at RT in 1X Exonuclease buffer containing 0.1 U/μL of Exonuclease III (EN0191). Secondary antibodies were added as described above.

### Image acquisition and analyses

#### Organoids

Images were acquired using a MICA Leica microscope with either a 43x oil immersion objective or a 63x water immersion objective, depending on the experimental requirements. The microscope settings were adjusted accordingly to optimize resolution and image quality for each magnification. For quantification of GFP^+^ CRISPRi cells in electroporated ventricles, all GFP^+^SOX2^+^, GFP^+^NEUROD2^+^, and GFP^+^TBR2^+^ cells were counted, and double positive cells were normalized to the total number of electroporated GFP^+^ cells.

#### Mice

Control and hJUNB samples were imaged at the confocal microscope LSM800. For embryonic stages, three sections per embryo were imaged and analysis was focused only on EdU^+^GFP^+^ cells with SOX2 and/or EOMES positive signal after image acquisition. SOX2^+^EOMES^-^ cells in the VZ were considered as RGs, EOMES^+^ cells in the VZ/SVZ were considered as IPs, and SOX2^-^ cells in the VZ/SVZ and EOMES^+^ cells in the CP were considered as neurons. At P1, 3-4 sections per pup were imaged and analyzed and values were averaged per animal: NEUROD2^+^BrdU^+^GFP^+^ neurons in the cortical plate were quantified and compared to all NEUROD2^+^GFP^+^ cells. Due to sparsity of the FUCCI vector electroporation, only one large region with at least 5 z stacks were imaged per biological replicate. All FUCCI positive cells in the VZ/SVZ were counted per image. mAG^+^mCherry^-^ cells were considered as S/G2/M, mAG^-^mCherry^+^ as G1 only and mAG^+^mCherry^+^ as G1/S transitioning cells. In all conditions, cells were hand counted using Fiji Cell Counter (76) in a blind manner, and proportional values were compiled in Excel and Graphpad Prism 10.

### Statistics

All statistical analyses were performed using Graphpad Prism 10.

#### Organoids cell type proportions

As the data did not follow a normal distribution, Mann-Whitney test was used to compare the RG / N ratio between control and JUNB KO electroporated cells.

##### Mouse *in utero* electroporations

###### Embryonic cell type proportions

Unpaired two-tailed t-test was used to compare the RG / IP-N ratio between control and JUNB electroporated cells, after passing Shapiro-Wilcoxon normality test.

###### Birthdating

Unpaired two-tailed t-test was performed to compare the proportions of BrdU positive control and JUNB electroporated neurons in the CP, after passing Shapiro-Wilcoxon normality test.

###### Clone size

Chi2 test was performed to compare control and JUNB clone size distributions.

###### Cell cycle phases

Non-parametric Mann-Whitney test was performed per cell cycle phase measurement, *i.e.* G1, G1/S, S/G2/M.

### Collection and sn/scRNA-seq of electroporated cells

#### E16.5 dataset

Twenty-four hours following electroporation, pregnant female was anaesthetized and sacrificed. Embryos from the pregnant mouse were harvested. Embryos were first separated based on electroporated plasmids (*i.e.* Control *vs.* JUNB), decapitated and the cortex was dissected out in 1X HBSS. The GFP^+^ cortices were separated and the electroporated region was dissected and frozen straight away in 2 ml Eppendorf tubes. Four cortices were collected per condition.

On the day of the snRNA-seq, samples were first gently thawed on ice and resuspended in 100 ul of lysis buffer composed of: 10 mM Tris-HCl (pH 7.4), 10 mM NaCl, 3 mM MgCl_2_, 0.01% NP-40, 1% BSA and 1 mM DTT. After 5 minutes of lysis on ice, 1 mL of HBSS was added. The samples were then centrifuged at 500 g for 5 minutes. After removing supernatant, the pellet was resuspended in 1 mL washing buffer composed of: 10 mM Tris-HCl (pH 7.4), 10 mM NaCl, 3 mM MgCl_2_, 1% BSA, 1 mM DTT and 0.5 U/μl RNAase inhibitor. Then, using the P1000 pipette, the samples were pipetted up and down 10 times. The samples were then centrifuged again for 5 minutes at 500 g. Supernatant was removed, 1 mL of additional washing buffer was added, and the pellet was resuspended. At this point, no clumps were seen, and the samples were passed through a 30-μm filter before FANS. Hoechst was used as a positive nuclei marker and 25’000 events representing Hoechst^+^GFP^+^ nuclei were sorted directly into empty 1.5 mL Eppendorf-tubes using Beckman Coulter MoFlo Astrios. Exactly 42.8 μL of the sorted nuclei were loaded on the first step of the 10x Next GEM 3’ reagent kit. The rest of the protocol was followed according to the manufacturer’s guidelines. The cDNA libraries were quality controlled using Agilent 2100 Bioanalyzer and TapeStation. Sequencing was performed using the NovaSeq 5000 platform with a 28/90 read configuration.

#### P0 dataset

Four days after electroporations and after birth of pups, four litter-mates per condition were sacrificed by decapitation and the cortex was dissected out of the head in cold 1x HBSS in less than 5 mins. The electroporated region was microdissected in a fresh petri dish containing 1x HBSS and transferred into an eppendorf tube. Dissociations were performed using the Neural Dissociation kit (Miltenyi, 130-092-628) by treating the microdissected tissue with Enzyme mix 1 for 7mins and then Enzyme mix 2 for 3mins. Cells were washed in sorting buffer (1x PBS, 1% BSA and 0.5 U/μl SUPERase-In RNase inhibitor (ThermoFisher Scientific, AM2694)) twice at 4 C with 300 g. 16,000 for Control and 50,000 Junb electroporated RFP^+^ cells were sorted into separate tubes containing sorting buffer. Sorted cell suspension was centrifuged again for 5 mins at 4 C with 300 g and all but around 43.2 μL was removed. The cell suspension was resuspended gently and processed with 10X Genomics Chromium Single Cell 3’ v3 reagent kit on the Chromium X controller. cDNA was split to generate a gene expression and a lineage barcode library as previously described (51). The quality control of the cDNA and libraries was performed using Agilent’s 2100 Bioanalyzer. Sequencing was performed using the NovaSeq 5000 platform with a 28/90 read configuration, aiming for a depth of 20,000 reads/cell for gene expression library and 2000 reads/cell for lineage barcode library.

### sn/scRNA-seq data analyses

#### Data preprocessing and quality controls

Fastq files were mapped on the standard 10X mouse reference genome GRCm38 using cellranger (77) (version: 7.2.0) for the E16 collected snRNA-seq dataset. For the P0 collected scRNA-seq, fastq files were mapped with the mouse reference genome including PBase, TurboRFP-NLS and hJUNB sequences for quality control purposes. Counts due to ambient RNA and random barcode swapping were removed from the raw matrix using remove-background function of cellbender (78) (version: 0.3.0) with epochs = 150, expected-cells=2000 for the E16 snRNA-seq dataset. The newly generated snRNA-seq cellbender count matrix was used for subsequent analysis. scRNA-seq P0 collected dataset was not processed through cellbender. Briefly, for the E16 collected dataseet, cells with nCount_RNA < 25’000, nFeature_RNA > 1’500 and percent.mt < 0.25 were included. For the P0 collected dataset, nCount_RNA < 30’000, nFeature_RNA > 1’000 and percent.mt < 10 were included. scDblFinder package (79) (version: 1.16.0) was used to remove doublets by simulating ncol(sce)/1000*0.008 doublet ratio.

#### Cluster definition and annotation

We used Seurat pipeline (80) (version: 5.0.1) to normalize and scale the data, and performed PCA analysis using the 3’000 most variable features. We performed umap dimension reduction using 30 PCs and use the FindNeighbors and FindClusters (resolution = 0.8) functions to define clusters, which we annotated based on their expression of marker genes *Sox2* and *Pax6* for RG, *Eomes* and *Neurog2* for IP and *Neurod2* and *Neurod6* for neurons.

#### Cell cycle scoring

We used the CellCycleScoring function (81) from Seurat package to define the cell cycle phase of our progenitor populations. Expression of RG human/primate specific genes in control / JUNB RG at different cell cycle phase was assessed using DotPlot.

#### Differential gene expression

We used the FindMarkers function from Seurat package (80) with min.pct = 0.2 to find genes differentially expressed between control and JUNB conditions.

#### Lineage barcode analysis

Lineage barcode library fastq files were processed as previously described (51) with nReads=10, nUMI=4 and nHamming=3 for both datasets. Control and Junb datasets were processed separately and merged after cloneID assignment. Interneuron containing clones were removed and the two datasets were balanced to have an equal number of cells for analysis.

### Preprocessing of human multiome dataset and base GRN construction

To perform *in silico* perturbation simulations, the published multiome dataset on Pcw21 developing cortical samples was analysed from GSE162170(31). Using scanpy (version 1.9.3), an anndata object was generated using available Multiome RNA counts, metadata and cluster names in .txt format. As the varnames were in ensembl_id, it was first to gene_symbol format. The anndata object with unique gene symbols was saved and pre-processed by filtering cells based on max_counts=20000 and min_counts=1500. Genes were filtered with min_counts=1 and the dataset was normalized the dataset with ‘RNA.counts’. Using canonical markers and rank_genes_groups function in scanpy, RG, IP, and N in the dataset were annotted and the rest of the celltypes were subsetted out. Cellcycle scoring was performed and top 3000 differentially expressed genes in the anndata were filtered. UMAP, PAGA, and ForceAtlas2 graph construction was performed to visualize the three broad celltypes. To end this step, one random cell in the RG population was selected as the root_cell to perform pseudotime calculation using CellOracle (version 0.10.15) and anndata object was saved with the calculated pseudotime.

Consensus ATAC peaks in .txt format from the same human multiome was processed to generate a bed file denoting the coordinate location of each ATAC peak. Peaks at the transcription start site (TSS) of genes were annotated and this processed peak file was saved in .csv format. Next, CellOracle wrapper for gimmemotifs (version 0.17.0) was used to generate a base GRN containing motifs in peaks close to 2kb from all gene promoters. Base GRN containing ‘motifs to peak coordinates’ and the final ‘TF to peak coordinates’ dataframes were saved as .csv files. These were used for motif analysis later.

### Motif analysis of JUNB promoter in mouse and human

#### Mouse

Consensus ATAC peaks from published mouse Multiome dataset (GSE241429) generated from E14, E15, and E16 developing cortical samples was used. Same as the human dataset, CellOracle wrapper for gimmotifs was used to generate a base GRN and final .csv files were either linking ‘motif to peak coordinates’ or ‘TF to peak coordinates’ were generated.

#### Human

The ‘motif to peak coordinates’ and ‘TF to peak coordinates’ .csv files generated from Human multiome dataset were used to generate a .csv file per gene.

For both datasets, the ‘motif to peak coordinates’ and ‘TF to peak coordinates’ .csv files contained all motifs detected by gimmemotifs at the peaks close to the promoter (<2kb) with an enrichment score. These .csv files were used to plot motif enrichment scores per gene per species.

### *In silico* perturbation of human JUNB using CellOracle

Once the RNA and ATAC from the published dataset was pre-processed, GRN construction using the CellOracle (version 0.10.15) package was performed through known TF to gene interactions from online databases(82) (TFLink). To predict dynamics states of expression, imputation of top 3000 expressed genes was performed according to the CellOracle tutorial, and these imputed values were used for *in silico* perturbations. A .csv file was generated for each celltype specific GRN. Finally, *in silico* KO (0 value of imputed_counts) and OE (maximum value of imputed_counts) was performed. A custom color scheme was used to guide the reader with the direction of the arrows following the perturbation.

## Code and data availability

All landscapes of gene expression and correlation values are available on https://www.humous.org/. Code used for the CellOracle, GRN and lineage barcode analysis was deposited on https://github.com/awaisj14/Humous_CellOracle/. All raw and processed single cell transcriptomics data generated in this study will be submitted to GEO.

## Funding

The Jabaudon laboratory is supported by the Swiss National Science Foundation, the Carigest Foundation, the European Research Council, the NeuroNA Foundation, the Roger de Spoelberch Foundation, the ERA-NET NEURON JTC, the Victoria Fund King Baudouin Foundation and the MCHRI Uytengsu-Hamilton 22q11 Neuropsychiatry Research Program.

## Author contributions

Conceptualization: E.K. & D.J.

Methodology: E.K., D.J., L.G., A.J., V.P., S.C.

Investigation: E.K., L.G., A. J., V.P., M.S., Q.LG.

Visualization: E.K., D.J., L.G., A. J., V.P.

Funding acquisition: D.J., S.C.

Supervision: E.K., D.J. & S.C.

Writing – original draft: E.K. & D.J.

Writing – review & editing: E.K., D.J., A.J. with the help from all authors.

## Acknowledgements

We thank the IGE3 genomics, Bioimaging and FACS platforms at University of Geneva for technical support. We also thank the FACS HCNP platform at Campus Biotech. We thank A. Benoit for technical assistance, L. Frangeul for logistical support, and all the members of the Jabaudon and Klingler labs for feedback on the manuscript. We thank UMANOVA (https://umanova.com) for developing the www.humous.org website.

## Competing interests

The authors declare no competing interests.

## Notes

https://www.humous.org/

### Competing Interest Statement

The authors have declared no competing interest.

