## Supplementary Figures for "Developmental gene expression patterns driving species-specific cortical features"

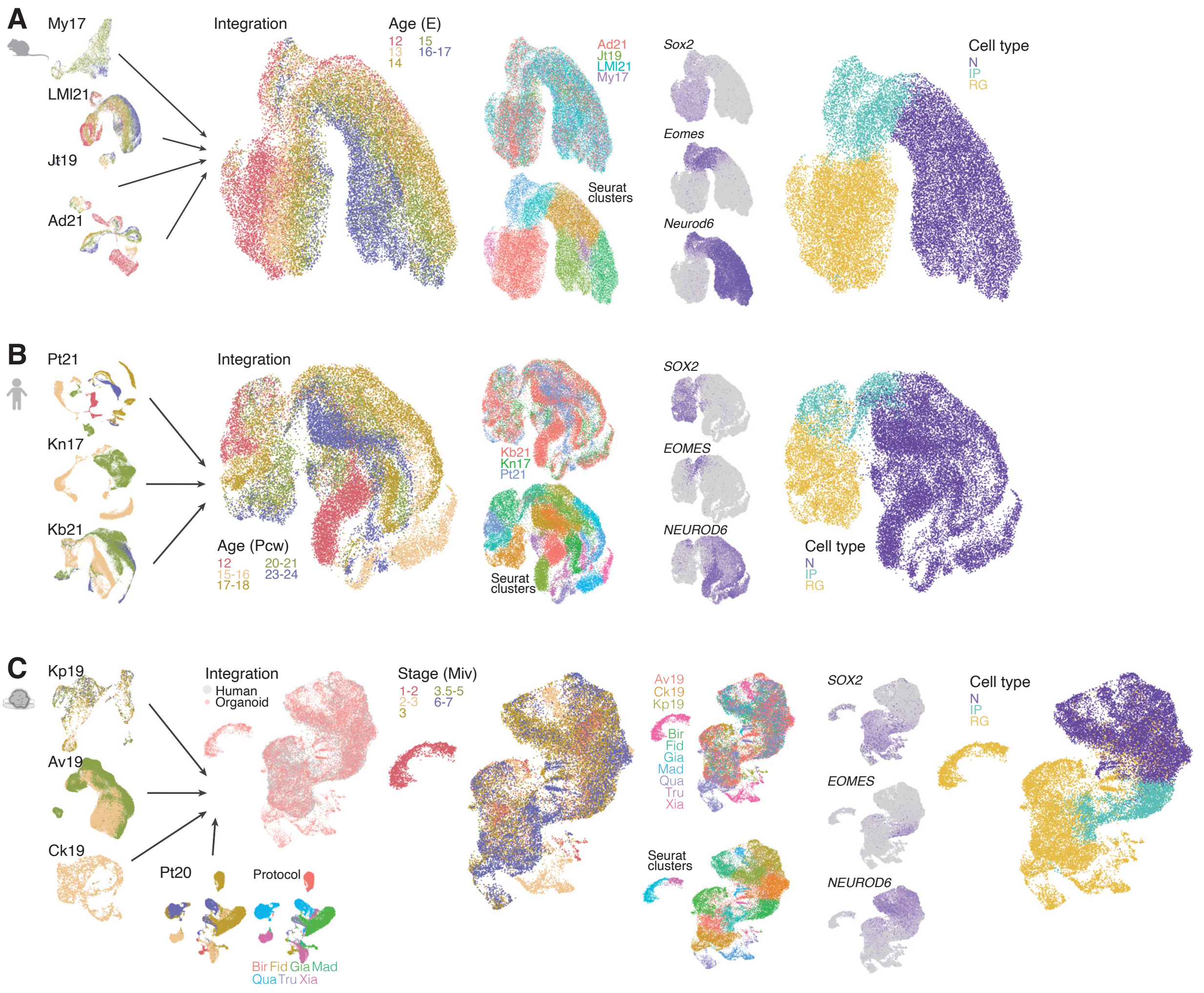


**Supplementary fig. 1. Integration of single-cell RNA sequencing datasets and cell type annotation consensus. (A-C)** Integration of single-cell RNA sequencing datasets from mouse (A) and human (B) developing cortex, and human-derived cortical organoid (C) (see **Table 1**). From left to right: individual dataset UMAPs annotated by age; integrated Mo, Hu, and $Org_H$ datasets (see **methods**) labeled by age, dataset (top right), and seurat clusters (bottom right); feature plots showing expression of *SOX2*, *EOMES* and *NEUROD6*; cell type annotation consensus. Abbreviations: Pcw, postconception week; E, embryonic day; Miv, month in vitro; RG, radial glia; IP, intermediate progenitor; N, neuron.

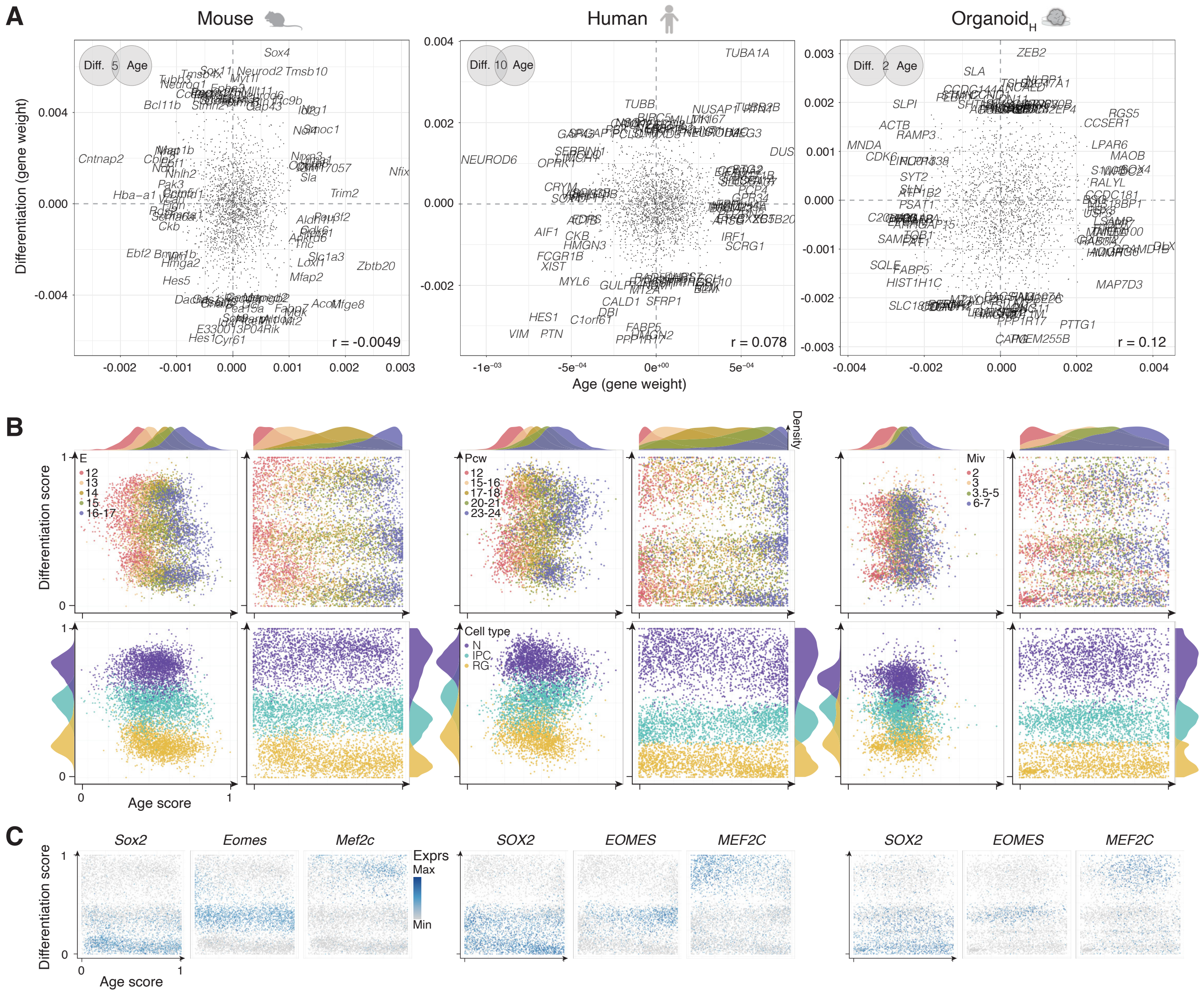


**Supplementary fig. 2. Differentiation and age prediction scores using ordinal regression models.**
**(A)** Top gene weights in the age and differentiation mouse (left), human (middle), and Org$_H$ (right) models. Number of common genes in the differentiation and age models are indicated in the venn diagrams for each condition. Correlation between age and differentiation weights (r) is indicated at the bottom right of each plot. Right: Common top genes between conditions in the differentiation (top) and age (bottom) models.
**(B)** Single cell prediction scores for age (x axis) and differentiation (y axis) before (left) and after (right) normalization for each condition. Cells are labeled by age (top) and differentiation (bottom). Densities along age and differentiation axes are shown for each condition.
**(C)** Expression of *SOX2*, *EOMES* and *MEF2C* in single cells along age and differentiation axes for each condition. Abbreviations: Diff., differentiation; Pcw, postconception week; E, embryonic day; Miv, month in vitro; Exprs, expression.

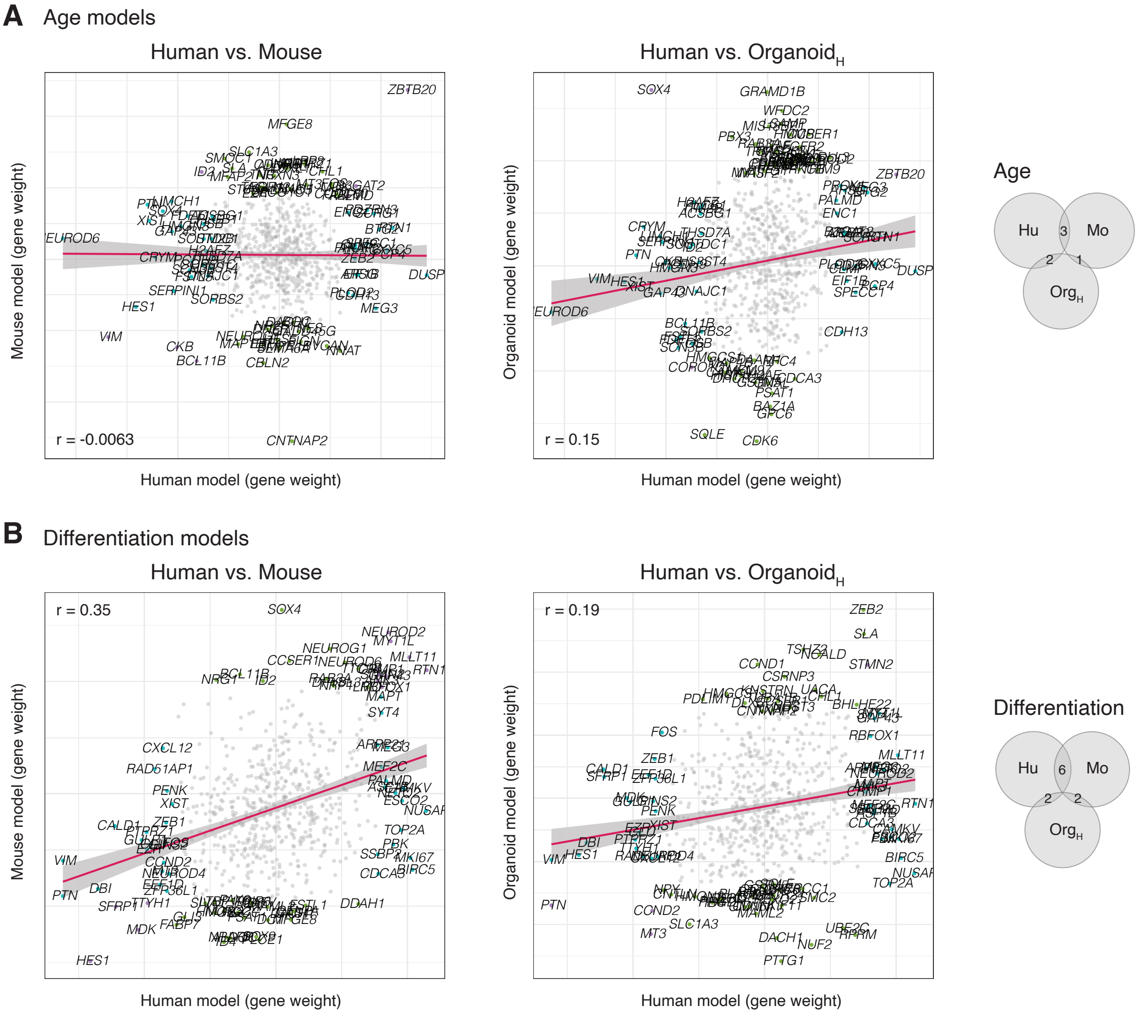


**Supplementary fig. 3. Comparison of gene weights between conditions in the age and differentiation models.** **(A)** Gene weight comparisons in human vs. mouse (left) and human vs. organoid$_H$ (right) age models. Correlations between gene weights are indicated (r). Right: Common top genes between conditions.
**(B)** Gene weight comparisons in human vs. mouse (left) and human vs. organoid$_H$ (right) differentiation models. Correlations between gene weights are indicated (r). Right: Common top genes between conditions.

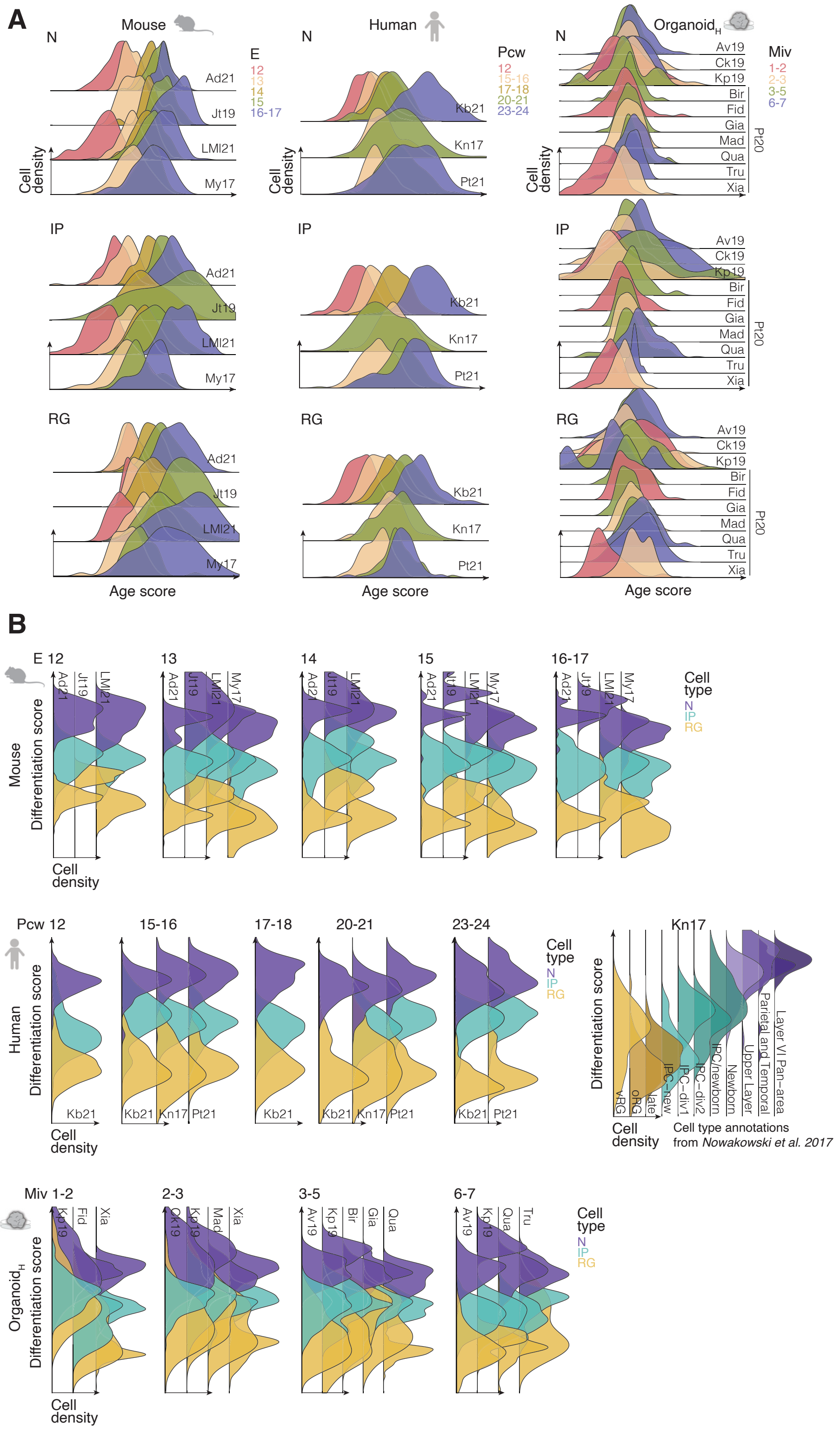


**Supplementary fig. 4. Differentiation and age predictions by dataset, age, and cell type.**

**(A)** Density of cells along the age prediction axis by their age at collection, for each dataset (see **Table 1**) and cell type (top, N; middle, IP; bottom, RG).

**(B)** Density of cells along the differentiation prediction axis by cell type annotation, for each dataset and age at collection (top, N; middle, IP; bottom, RG).

Abbreviations: Pcw, postconception week; E, embryonic day; Miv, month in vitro; N, neuron; IP, intermediate progenitor; RG, radial glia.

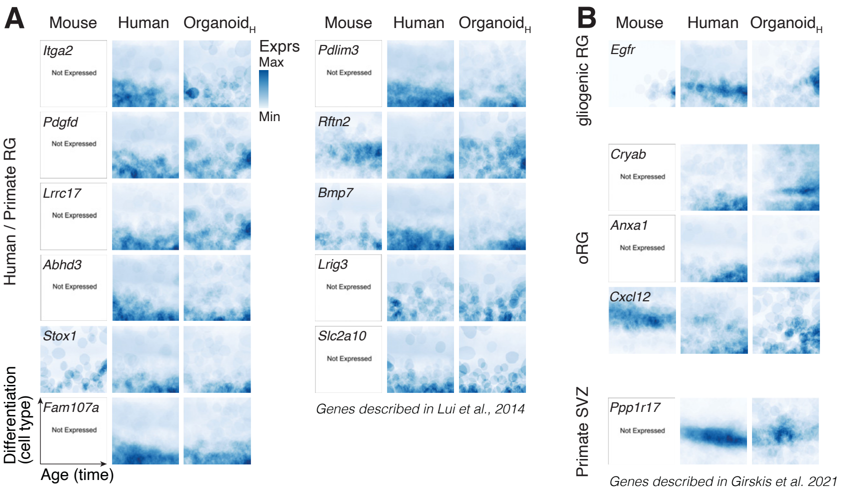


**Supplementary fig. 5. Human/primate radial glia-specific markers in Humous.org browser.**
Expression landscapes of genes shown to be specific to human/primate RG **(A)**, gliogenic RG and oRG **(B, top)** in Lui et al., 2014 (85) and to primate SVZ **(B, bottom)** in Girskis et al., 2021 (83).
Abbreviations: RG, radial glia; oRG, outer RG; SVZ, subventricular zone.

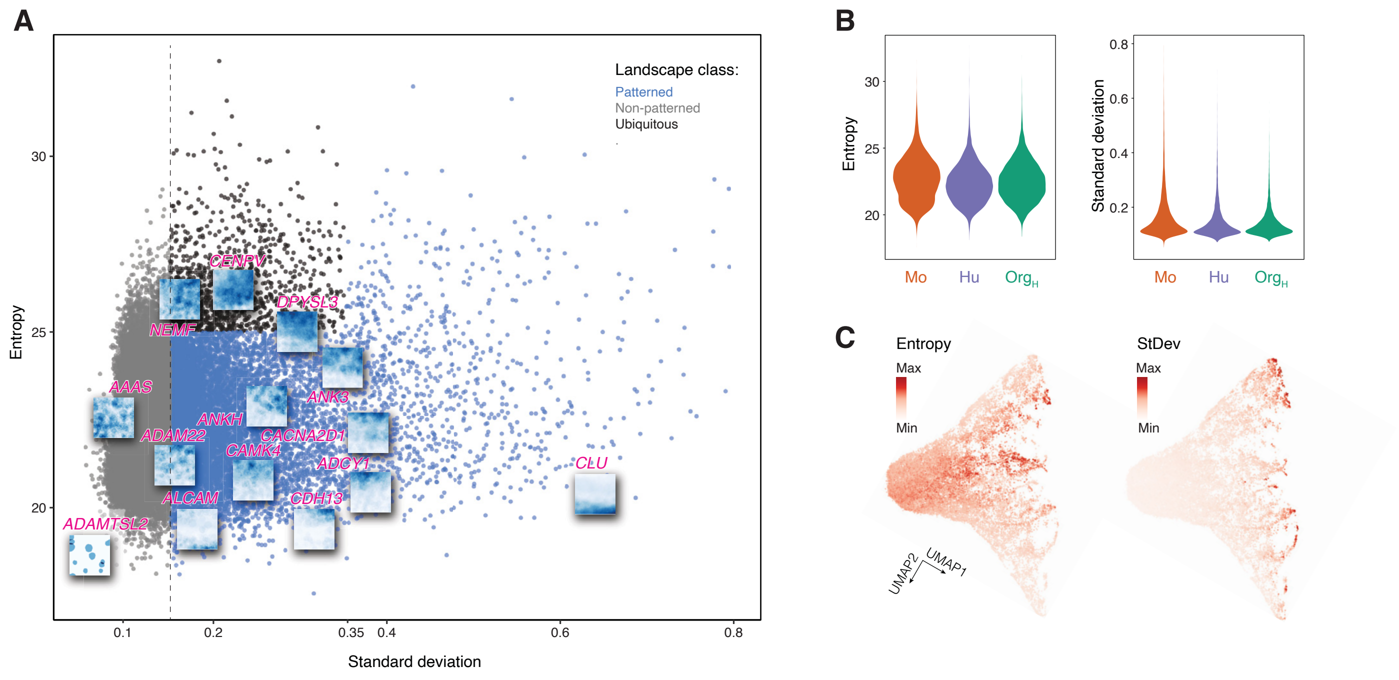


**Supplementary fig. 6. Entropy and standard deviation define classes of gene expression landscapes.**
**(A)** Entropy and standard deviation of all landscapes and definition of landscape classes. Representative examples of human gene expression landscapes are shown.
**(B)** Entropy (left) and standard deviation (right) distributions of all landscapes in Mo, Hu, and $Org_H$.
**(C)** Entropy (left) and standard deviation (StDev, right) of all landscapes in the umap space.

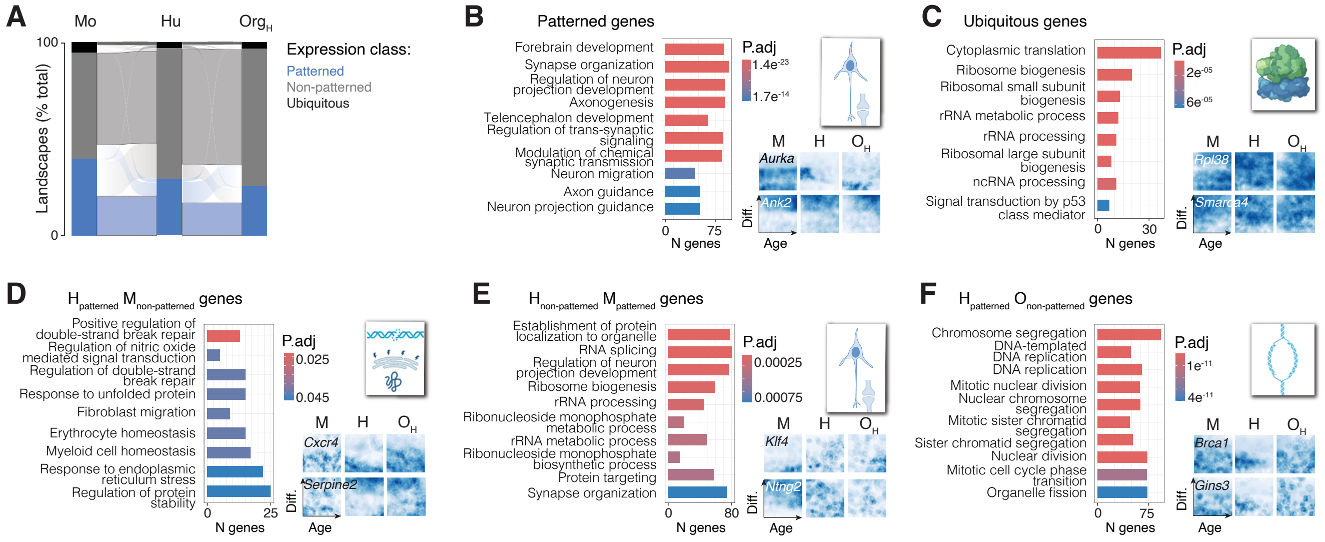


**Supplementary fig. 7. Ontologies of genes with specific expression classes across conditions.**
**(A)** Comparison of landscape classes between conditions for corresponding genes.
**(B-C)** Top 10 ontologies of genes with patterned (B) and ubiquitous (C) expression landscapes in all conditions.
**(D)** Top 10 ontologies of genes with patterned expression landscapes in Hu ($H_{patterned}$) which are non-patterned in Mo ($M_{non\text{-}patterned}$).
**(E)** Top 10 ontologies of genes with non-patterned expression landscapes in Hu ($H_{non\text{-}patterned}$) which are patterned in Mo ($M_{patterned}$).
**(F)** Top 10 ontologies of genes with patterned expression landscapes in Hu ($H_{patterned}$) which are non-patterned in $Org_H$ ($O_{non\text{-}patterned}$).
In B-F, Insets summarize the main gene ontologies (top right) and examples of gene expression landscapes are shown for each conditions (bottom right). Only significant gene ontologies are shown.
Abbreviations: p.adj, adjusted p-value; Diff., differentiation.

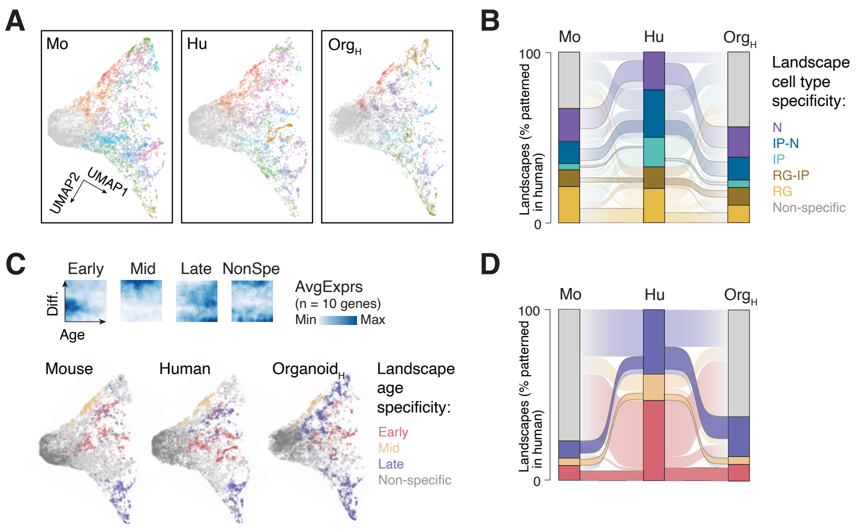


**Supplementary fig. 8. Cell type- and age-specific landscapes across conditions.**
**(A)** Landscapes clusers in Mo, Hu, and OrgH labeled in the umap space.
**(B)** Corresponding landscape cell type specificity across conditions.
**(C)** Early, mid, and late specific landscape clusters (top) and their distribution in the umap space (bottom).
**(D)** Corresponding landscape age specificity across conditions.
Abbreviations: N, neuron; IP, intermediate progenitor; RG, radial glia; NonSpe, non-specific; AvgExprs, average expression.

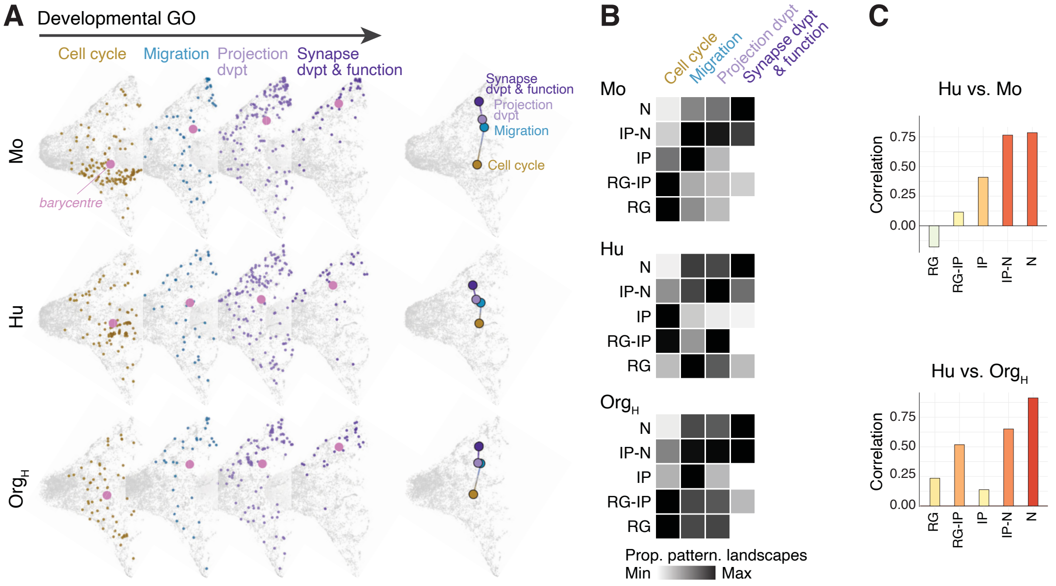


**Supplementary fig. 9. Cell type-specific landscapes of genes associated with developmental ontologies.**
**(A)** Landscapes of genes associated with cell cycle, migration, projection development and synapse development and fonction ontologies in Mo, Hu, and $Org_H$ (left), and their barycentre for each ontology (right).
**(B)** Landscapes cell type specificity across developmental gene ontologies in Mo, Hu, and $Org_H$.
**(C)** Correlations of gene ontologies for landscapes with corresponding cell type specificity between species (top) or contexts (bottom).
Abbreviations: N, neuron; IP, intermediate progenitor; RG, radial glia; GO, gene ontology; dvpt, development.

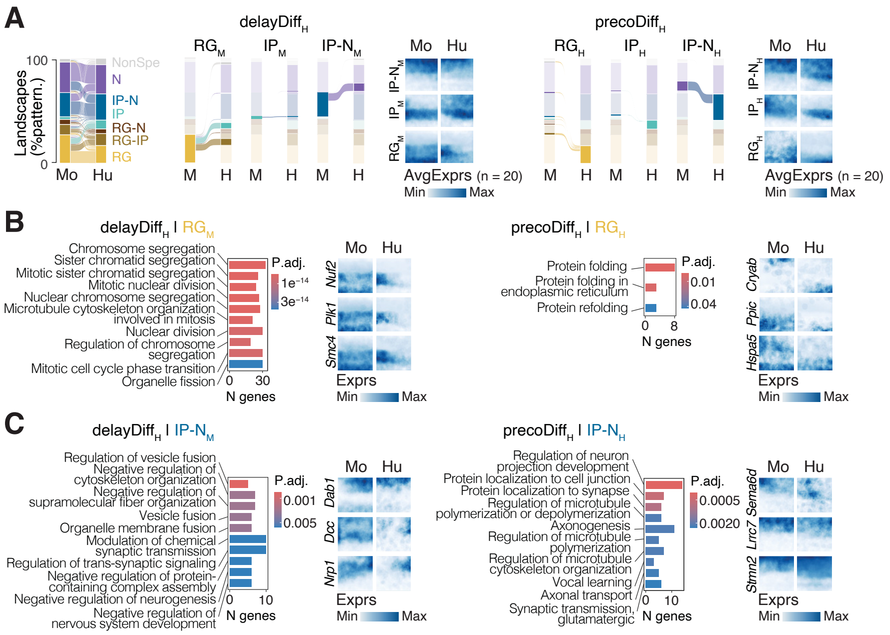


**Supplementary fig. 10. Delayed and precocious human gene expression profiles.**
**(A)** Left: Corresponding landscape cell type specificity between species. Middle: Definition of landscapes with delayed differentiation specificity in Hu (delayDiff$_H$) and their average expression (AvgExprs) in Mo and Hu. Right: Definition of landscapes with precocious differentiation specificity in Hu (precoDiff$_H$) and their average expression in Mo and Hu.
**(B)** Left: Top 10 ontologies of delayDiff$_H$ genes expressed in Mo RG. Right: Top 10 ontologies of precoDiff$_H$ genes expressed in Hu RG.
**(C)** Left: Top 10 ontologies of delayDiff$_H$ genes expressed in Mo IP-N. Right: Top 10 ontologies of precoDiff$_H$ genes expressed in Hu IP-N.
In B-C, Examples of gene expression landscapes related to the ontologies are shown in Mo and Hu. Only significant gene ontologies are shown.
Abbreviations: N, neuron; IP, intermediate progenitor; RG, radial glia.

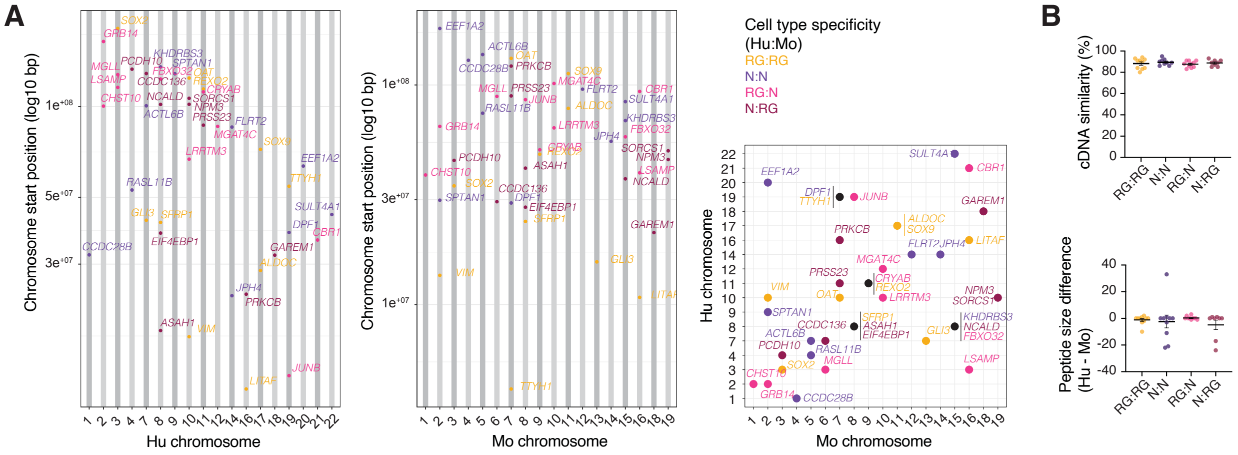


**Supplementary fig. 11. Genomic and protein features of genes with convergent or divergent expression landscapes between species.**

**(A)** Hu (left) and Mo (middle) chromosome position of genes expressed in RG (yellow) or N (purple) in both Mo and Hu (convergent expression), genes expressed in Mo RG and Hu N (brown), and genes expressed in Mo N and Hu RG (magenta) (divergent expression). Right: Comparison of chromosome positions in Hu and Mo.

**(B)** Hu and Mo cDNA similarity and encoded peptide size difference of genes with convergent and divergent expression.

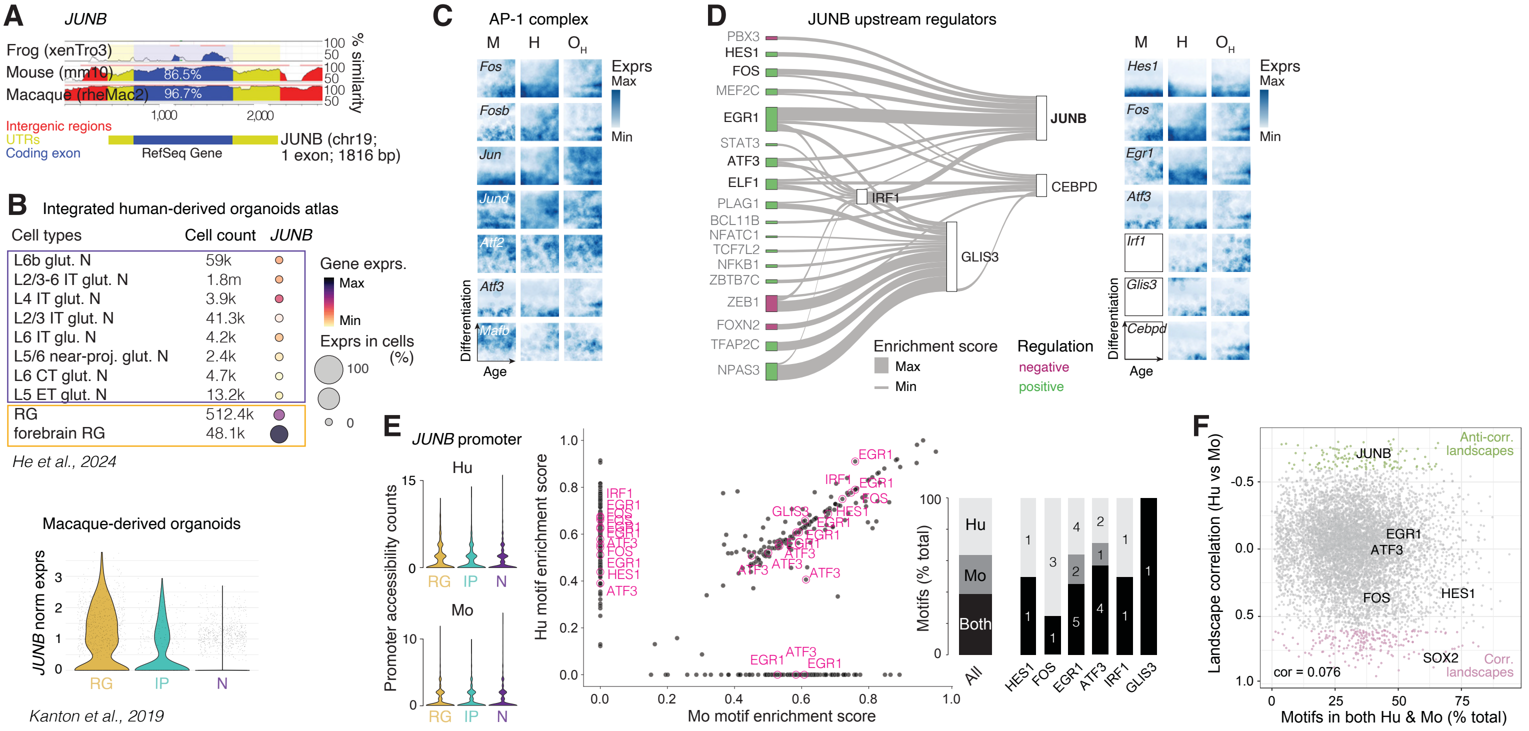


**Supplementary fig. 12. Specific expression of JUNB in human / primate radial glia and upstream regulators.** **(A)** Similarity in *JUNB* coding exon, UTRs and intergenic regions in Frog, Mouse, and Macaque compared to Human. Data are from the UCSC Genome Browser (Perez et al., 2025 (84)).

**(B)** Expression of *JUNB* in human-derived (from He et al., 2024 (39)) (top) and macaque-derived (bottom) organoids (Kanton et al., 2019 (48)) single-cell RNA sequencing data. Cell type annotations are from the original publications.

**(C)** Expression landscapes of genes coding for other AP-1 complex members in Mo, Hu, and $Org_H$.

**(D)** Left: Sankey plot displaying predicted upstream regulators of *JUNB*, *CEBPD*, *IRF1* and *GLIS3*. The line width connecting upstream regulators (left) to downstream targets (right) reflects mean coefficient score calculated from the CellOracle package. The predicted positive (green) and negative regulation (magenta) is determined by the sign of mean coefficient score. Factors expressed in RG are in bold. Right: Expression landscape examples of *JUNB* RG-specific upstream regulators. *Irf1*, *Glis3* and *Cebpd* genes were filtered out from Mo dataset because of too low expression.

**(E)** Left: Hu and Mo *JUNB* promoter accessibility counts per cell type. Center: Hu and Mo motif enrichment scores in *JUNB* promoter. Right: Proportion of *JUNB* promoter motifs present in both species, Mo only, or Hu only for all motifs (all), or motifs associated with Hu RG-specific transcription factors.

**(F)** Correlation between landscape similarity between species and proportion of motifs present in both species.

Abbreviations: RG, radial glia; IP, intermediate progenitor; N, neuron; Exprs, expression.

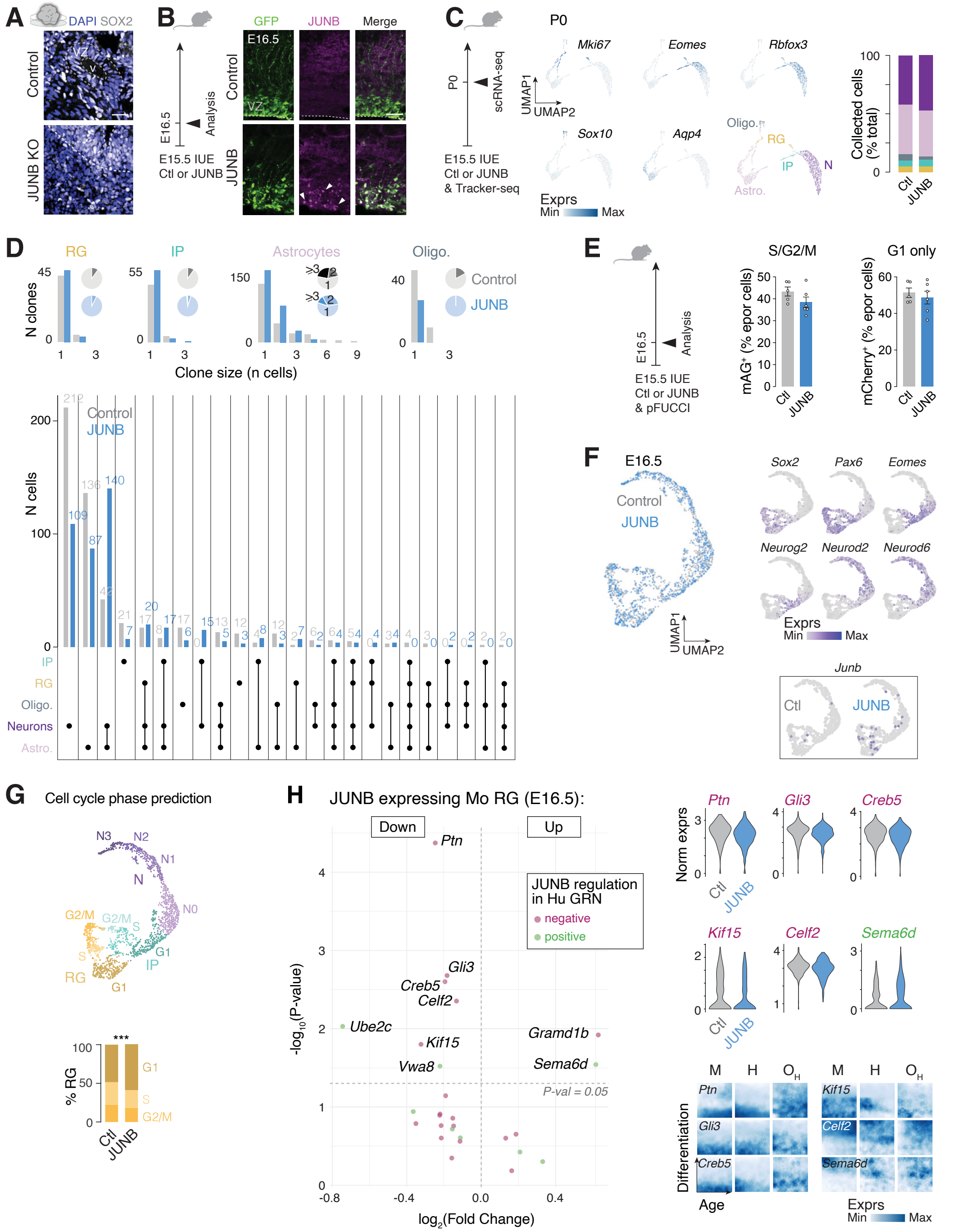


**Supplementary fig. 13. JUNB expression changes mouse radial glia properties and neuronal output.**

**(A)** Photomicrographs illustrating how ventricles and VZ are defined in organoids based on SOX2 and DAPI stainings.

**(B)** JUNB protein in Mo VZ electroporated cells 24 hours after in utero electroporation with pCS2::JUNB plasmid.

**(C)** Tracker-seq experiment. Left: Feature plots with marker genes and cell type definition in the umap space. Right: Proportion of cells in control and JUNB conditions.

**(D)** Top: Clone size by cell types. Bottom: Number of cells per clone composition found in the Tracker-seq data.

**(E)** Quantifications of electroporated RG in S/G2/M or G1-only phases using pFUCCI.

**(F)** Single-nucleus RNA-sequencing data of control and JUNB in utero electroporated cells at E15.5, collected at E16.5. Left: Control and JUNB cells in the umap space. Middle: marker genes used for cell type annotations. Right: *Junb* expression in control and JUNB conditions.

**(G)** Top: Cell cycle phase prediction by cell type. Bottom: Proportions of RG in G1, S and G2/M phase in control and JUNB conditions.

**(H)** Left: Genes of Hu JUNB gene regulatory network (GRN) differentially expressed between control and JUNB Mo RG. Only genes with p-value (p-val) < 0.05 are labeled. Right: Expression of genes with differential expression in Mo RG, which mirrors JUNB predicted regulation in Hu GRN. Top: violin plots showing expression in control and JUNB Mo RG. Bottom: Expression landscapes in Mo, Hu, and OrgH.

Abbreviations: Ctl, control; N, neuron; IP, intermediate progenitor; RG, radial glia. Exprs, expression.
